# Single amino acid-promoted reactions link a non-enzymatic chemical network to the early evolution of enzymatic pentose phosphate pathway

**DOI:** 10.1101/2020.08.11.245860

**Authors:** Gabriel Piedrafita, Sreejith Varma, Cecilia Castro, Christoph Messner, Lukasz Szyrwiel, Julian Griffin, Markus Ralser

**Author notes:** Spanish National Cancer Research Centre (CNIO), Madrid, Spain.

## Abstract

How metabolic pathways emerged in early evolution remains largely unknown. Recently discovered chemical networks driven by iron and sulfur resemble reaction sequences found within glycolysis, gluconeogenesis, the oxidative and reductive Krebs cycle, the Wood Ljungdahl as well as the S-adenosylmethionine pathways, components of the core cellular metabolic network. These findings suggest that the evolution of central metabolism was primed by environmental chemical reactions, implying that non-enzymatic reaction networks served as a “template” in the evolution of enzymatic activities. We speculated that the turning point for this transition would depend on the catalytic properties of the simplest structural components of proteins, single amino acids. Here, we systematically combine constituents of Fe(II)-driven non-enzymatic reactions resembling glycolysis and pentose phosphate pathway (PPP), with single proteinogenic amino acids. Multiple reaction rates are enhanced by amino acids. In particular, cysteine is able to replace (and/or complement) the metal ion Fe(II) in driving the non-enzymatic formation of the RNA-backbone metabolite ribose 5-phosphate from 6-phosphogluconate, a rate-limiting reaction of the oxidative PPP. In the presence of both Fe(II) and cysteine, a complex is formed, enabling the non-enzymatic reaction to proceed at a wide range of temperatures. At mundane temperatures, this ‘minimal enzyme-like complex’ achieves a much higher specificity in the formation of ribose 5-phosphate than the Fe(II)-driven reaction at high temperatures. Hence, simple amino acids can accelerate key steps within metal-promoted metabolism-like chemical networks. Our results imply a stepwise scenario, in which environmental chemical networks served as primers in the early evolution of the metabolic network structure.

**Significance Statement:** The evolutionary roots of metabolic pathways are barely understood. Here we show results consistent with a stepwise scenario during the evolution of (enzymatic) metabolism, starting from non-enzymatic chemical networks. By systematic screening of metabolic-like reactivities *in vitro*, and using high-throughput analytical techniques, we identify an iron/cysteine complex to act as a ‘minimal enzyme-like complex’, which consists of a metal ion, an amino acid, and a sugar phosphate ligand. Integrated in a metal-driven, non-enzymatic pentose phosphate pathway, it promotes the formation of the RNA-backbone precursor ribose 5-phosphate at ambient temperature.

## Introduction

The evolutionary origins of enzymatic metabolism are subject to a long-lasting debate (1–6) but remain essentially unknown. The history of Biochemistry and the evolution of biochemical thinking have fundamentally shaped this debate. In earlier days, metabolism was seen primarily as the biological process of nutrient uptake and conversion, to provide the biomolecular building blocks that enable survival and growth of the cells. On the basis of this school of thought, the studies into the origin of metabolism during abiogenesis primarily focused on the nutritional aspects of metabolism, that is, to provide nucleotides, amino acids, lipids to early cells (7). Modern biochemistry has however strongly shifted to a view that the role metabolism exceeds the linear conception of its biosynthetic function (8, 9). Metabolism, and the biochemistry that operates within it, constraints the very existence of biological systems. Structured in a topologically conserved system (the metabolic network), metabolism determines the conditions under which cells can persist and how they can adapt and evolve in new environments, by controlling which elements and molecules are required for a cell to grow, how cells divide (e.g. by eliciting gene expression programs), which molecules are transported through their membrane, and managing chemical processes causing damage to cells (10–15). As a consequence of this paradigm shift in biochemistry, also research into the origins of metabolism is influenced. It has hence become important not only to consider scenarios for the origin of end-point metabolites, amino acids, nucleotides or lipids, as the sole presence of these molecules may not be sufficient to explain the necessary biological properties. Instead, we are urged to explain too the origin of the structure of the metabolic network (16–18).

With little experimental evidence available, the debate about the topological origin of metabolism was purely hypothetical, and split between an ‘evolutionary view’, in which the current structure of metabolism emerged over time as a consequence of the evolution of enzymes (4, 19, 20), and an environmental-chemistry view, that places non-enzymatic reactions at the root of the metabolic network structure (2, 3, 6). Indeed, an astonishing feature of the cellular metabolic network is the evolutionary conservation of its structure, basic function principles, and topology (21, 22). The universal conservation of these features indicates that the metabolic network dates back to the very early stage of evolution. This hypothesis is supported by the observation that the topological organization of metabolism is maintained much better than the (sequence) conservation of metabolic enzymes. An example of this is the glycolytic pathway, that is found in both Eubacteria and Archaea, but without a general conservation of the catalyzing enzymes (23).

More recently, direct experimental evidence that supports a non-enzymatic origin of the metabolic network structure has emerged. New sensitive analytical methods like multiple reaction monitoring have become available, and allow large screening experiments, and reach detection limits that are several orders of magnitude better compared to the method spectrum conventionally used in the origin of life research (24). Importantly, selective reaction monitoring experiments can be performed in the presence of ferrous iron concentrations that cause signal-suppression in 1H-NMR experiments that have frequently been used in origin of life research (25). Ferrous iron is believed to have reached millimolar concentrations in Archean oceans and other aquatic environments (26–28), is a biologically highly relevant catalyst and electron donor, and as catalyst of many metabolism-like non-enzymatic reactions, may be key in understanding early metabolic reactions (29, 30).

The ability to perform sensitive metabolite quantification experiments in Archean-like, iron-rich reaction conditions, led to the discovery of a series of chemical reactions that capture the topological organization of metabolic pathways, in a single condition and without an enzyme present. The first such network system resembles the Embden Meyerhoff Parnass pathway, or glycolysis, and the pentose phosphate pathway (PPP). This network is driven by the presence of Fe(II) concentrations that resemble those of average Archean sedimentation (29, 31). Meanwhile additional reaction systems have been described. These concern other glycolytic reactions, and suggest that at least gluconeogenesis, the oxidative as well as reductive Krebs cycle, its glyoxylate shunt, as well as the Wood-Ljungdahl pathway all have topologies that resemble those of simple non-enzymatic networks that could have occurred prior to the origins of life (30, 32–37). This ‘fuzzy’ chemistry hence captures major parts of central metabolism.

In this manuscript, we aim to address a question that emerges from these observations: How could a metal-ion driven, chemical network prime a Darwinian process that eventually forms an enzyme-catalyzed metabolic pathway? Indeed, as early as in 1945, Horowitz described a main difficulty to explain the advent of enzymatic catalysis by Darwinian evolution, as a chicken-egg dilemma, that we have previously referred to as the ‘End Product Problem’ (5, 38): Darwinian pressure can only select for a functional product, the ‘end product’ of an enzymatic pathway. But how could then evolve the enzyme that catalyzes the formation of an upstream intermediate, one that is needed before formation of the functional product is conceived?

We speculate that the presence of a non-enzymatic chemical network could overcome the end product problem (5). If a reaction network is already present due to non-enzymatic interconversion of metabolites, Darwinian selection can then improve the end product formation by selecting an enzyme that enhances its most rate-limiting reaction. In this model, not all enzymes need to come into place at once, but an improvement of the entire pathway is possible with the selection of a single enzyme (**Fig. 1A**). This model generates at least two predictions. The first is that in this model reaction topologies will not substantially change over time. Our biological knowledge is consistent with this prediction; the metabolic reaction topologies are strongly conserved among species and barely changed over billions of years of evolution (21, 22)). The second prediction is that in order to enable the stepwise evolution of enzymes, the simple components of enzymes (amino acids or small peptides), must be able to improve rate-limiting reactions of extant metabolic pathways in an enzyme-free setup.

**Figure 1.**
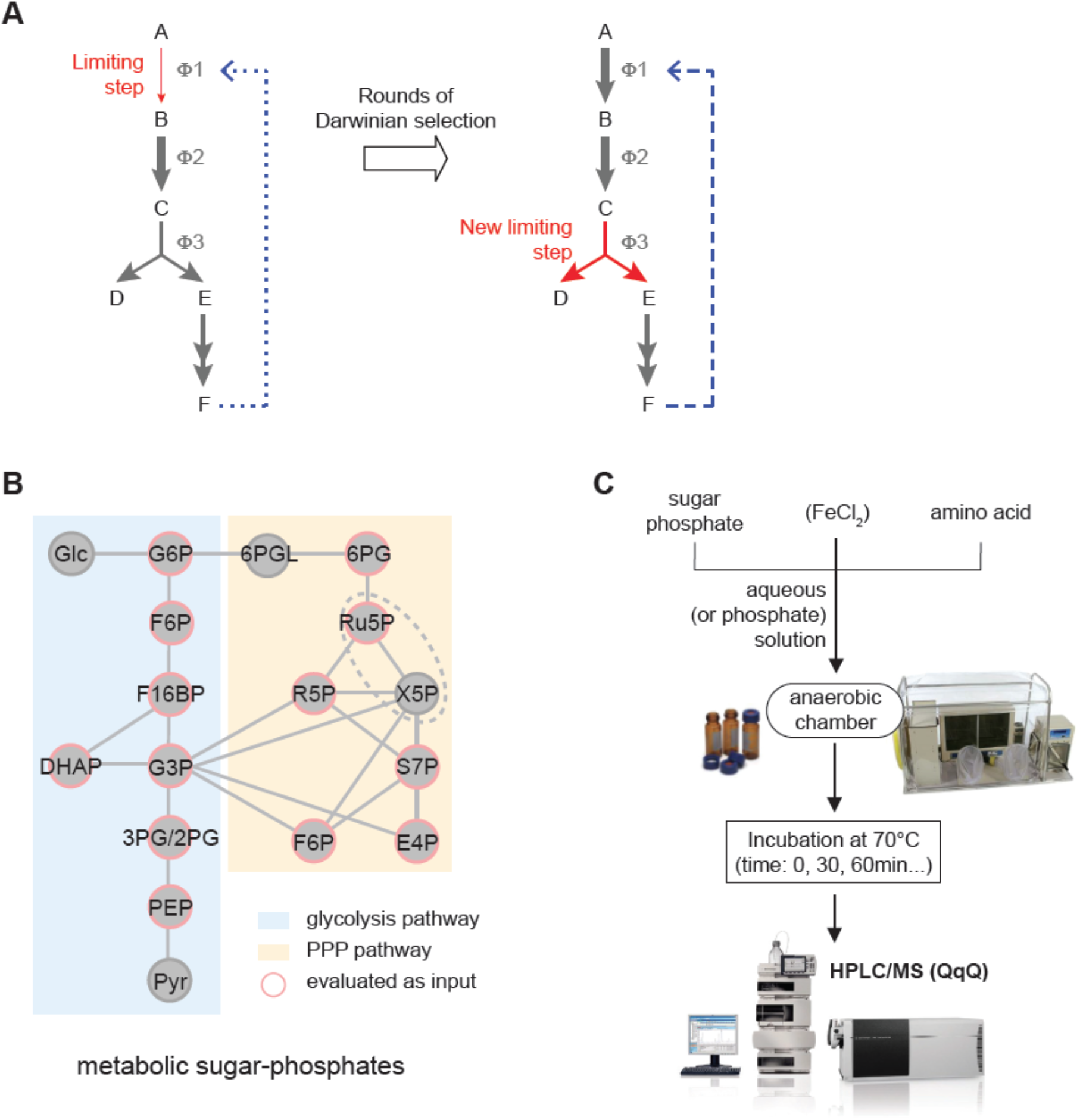
Model for a stepwise scenario in the early evolution of catalyzed metabolic networks, and experimental approach for screening non-enzymatic conversions among metabolic sugar-phosphates influenced by amino acids. (A) Hypothetical scheme of how a metabolic pathway could evolve from an underlying non-enzymatic network. Selective pressure for higher pathway product yield falls on the rate-limiting step (ɸ1, red). This would be improved by the selection of a primitive enzyme, until another process (ɸ3) becomes limiting. Some immediate or far end-products might display catalytic potential helping to fix those catalytic activities by feedback. (B) Network topology of metabolic sugar-phosphate interconversions from modern enzymatic glycolysis and pentose phosphate pathways. Enzymes and co-substrates are omitted in this graph and only sugar phosphates analyzed in this study are displayed (those used as input appear surrounded). Abbreviations used: Glc, glucose; G6P, glucose 6-phosphate; F6P, fructose 6-phosphate; F16BP, fructose 1,6-bisphosphate; DHAP, dihydroxyacetone phosphate; G3P, glyceraldehyde 3-phosphate; 3PG/2PG, 3-phosphoglycerate & 2-phosphoglycerate; PEP, phosphoenolpyruvate; Pyr, pyruvate; 6PGL, 6-phosphogluconolactone; 6PG, 6-phosphogluconate; Ru5P, ribulose 5-phosphate; R5P, ribose 5-phosphate; X5P, xylulose 5-phosphate; S7P, sedoheptulose 7-phosphate; E4P, erythrose 4-phosphate. (C) Experimental procedure. Each sugar phosphate was dissolved in aqueous (or 50 mM phosphate) solution in the presence or absence of single amino acids and/or FeCl2. Samples were incubated at 70°C in glass vials sealed in an anaerobic chamber (picture courtesy of Coy Laboratory Products Inc.) before targeted detection and quantification of sugarphosphate products (from all those in (B)) using a tandem liquid chromatography-mass spectroscopy (LC-MS/MS) system operating in SRM mode (Agilent Technologies).

To test this prediction, we chose a non-enzymatic network of glycolytic and PPP-like reactions as a starting point, so far described as Fe(II)-driven chemistry (29, 31). We then systematically combined the metabolites from these sugar-phosphate interconversions with the 20 universal proteinogenic amino acids, and followed the ~200 possible carbohydrate interconversion reactions by quantitative selective reaction monitoring (**Fig. 1B-C**). We discover that amino acids have broad impact on non-enzymatic metabolism like reactions on the non-enzymatic reactions that resemble glycolysis and the PPP. Strikingly, we identify one case in which an amino acid replaces a metal ion and promotes the formation of the RNA-backbone precursor ribose 5-phosphate from 6-phosphogluconate, a rate-limiting reaction of the extant oxidative PPP. Mimicking an evolutionary scenario where biochemical residues would start to accrue on metal ion centers, we observe the formation of a ternary complex between iron, cysteine and the sugar phosphate, that inspires enzyme-resembling properties in the formation of the RNA-backbone metabolite. Our results hence support a theory for the stepwise origin of enzymatic pathways on the basis of fuzzy chemical networks that serve as templates in the origin of enzymatic metabolism.

## Results

### Amino acids impact non-enzymatic metabolism-like interconversions between metabolites of glycolysis and the pentose phosphate pathway

A high-throughput *in vitro* screening was devised in order to assess the influence of amino acids on the reactivity of central carbon metabolites (**Fig. 1B-C**). Twelve commercially available sugar-phosphate intermediates -- seven belonging to glycolysis (Embden Meyerhof; 4 of those shared with Entner-Doudoroff) and five to the pentose phosphate pathway (PPP) -were incubated, at a concentration of 100 μM, in different aqueous solutions containing proteinogenic amino acids. Amino acids were grouped in sets of two or three according to their chemical similarity, and added at a total concentration of 400 μM. We chose concentrations much lower than those typical of cells, as precursors of metabolic pathways more likely had lower concentrations than metabolites in modern cells and lower concentrations are used in origin of life studies that are based on one-reaction-at-a-time principles of organic chemistry. We hence choose a highly sensitive LC-SRM method that reliably detects the metabolites down to attomoles (31). Samples were prepared in an artificial nitrogen atmosphere to simulate low oxygen concentrations of the early Archean atmosphere prior to the great oxygenation events.

In the screening setting, mixtures were heated to 70°C, a compromise defined previously to achieve detectable non-enzymatic reaction rates while still being within the temperature range where many hyperthermophiles relying on classical Embden-Meyerhof or variants of it that preserve the same metabolic pathway structure are known to grow (39). To capture the dynamic behavior of the non-enzymatic network letting intermediate metabolites not only to form but also to react further, we sampled at multiple points over a total experimental time of 6 hrs. Sugar-phosphate products were detected and quantified by targeted tandem liquid chromatography-mass spectroscopy (LC-MS/MS), from which we reconstructed the kinetics of metabolite formation and consumption (**Fig. 2**; **Materials and Methods**). Overall, a total of twenty-three non-enzymatic reactions that resemble glycolysis and the PPP were detected (a similar number as in Keller et al, 2014). The presence of amino acids did however show noticeable effects on the reaction rates of non-enzymatic glycolysis and PPP-like reactions (**Fig. 2A**; **Fig. S1**). We observed both the acceleration and the deceleration of several non-enzymatic conversions, with typical rate fold changes in the range 0.25-4.0. Overall, thirteen reactions were significantly enhanced by at least one group of amino acids, when compared to a control case without amino acids (Mann-Whitney *U* test, α = 0.10).

**Figure 2.**
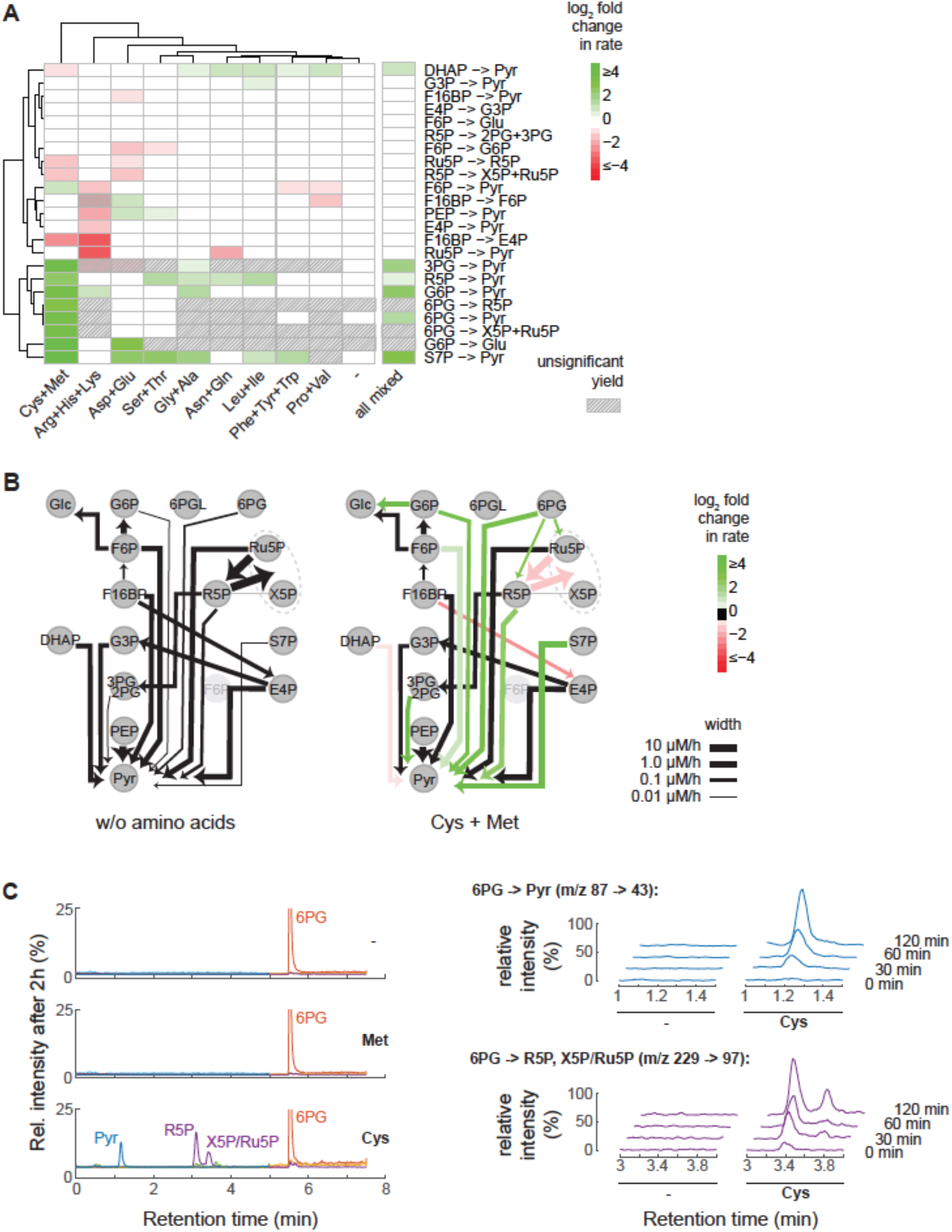
Amino acids impact non-enzymatic interconversions between glycolytic and pentose phosphate pathway sugar phosphates. A total of 12 metabolic intermediates from glycolysis and pentose phosphate pathway were incubated separately, at a concentration of 100 μM, at 70°C, in aqueous solutions containing single, proteinogenic amino acids (grouped, according to functional similarity, at a total concentration of 400 μM). The non-enzymatic formation of sugar-phosphate products was monitored over time by targeted LC/MS. (A) Amino-acid effects on individual production rates, shown in terms of log fold changes vs. a control case without amino acids, illustrated as a Heat-Map. Only individual transformations with over-background product detection and congruent increase between replicates in at least one amino-acid condition were considered in the analysis; striped cells denote particular conditions where these criteria were not fulfilled (i.e. insignificant product detected). Only cases of significant rate changes (Mann-Whitney *U* test, α = 0.1) are color coded. ‘all mixed’ corresponds with a condition where all 20 amino acids were present at a total concentration of 400 μM. *N* = 3 independent experiments per condition. (B) Graphs showing the non-enzymatic transformations detected in the aqueous control compared with those in the presence of cysteine and methionine. The width of the arrows is proportional to the calculated reaction rates (see Materials and Methods) and same color coding as in (A) is applied to transformations with significantly different rates (either higher - green - or lower - red) in the cysteine + methionine case vs. control (black = non-sig. change). (C) Representative chromatograms obtained by LC/MS after 2h incubation of 100 μM 6PG in control conditions without amino acids (-), in the presence of 200 μM methionine, or with 200 μM cysteine, revealing that cysteine is responsible for facilitating the formation of different products. Right panels: Detailed time courses in the formation of pyruvate and pentose phosphate sugars (SRM transitions displayed).

As one can expect from the nature of chemical reactions, a rate can be changed either by the direct interaction of an amino acid with the sugar phosphates, or by indirect effects, like changes in pH. A number of reactions were indeed changed by positively-charged amino acids (arginine, histidine and lysine) as well as by negatively-charged ones (aspartate and glutamate) and were hence possibly explained by their ability to change in response to shifts in the pH, an important factor in the non-enzymatic reactivity of sugar phosphates (Keller et al, 2016) (**Fig. S1**). In this work, however, we will focus on the most prominent changes observed with the sulfur-containing amino acids (cysteine and methionine) (**Fig. 2A-B**; **Fig. S1**). Eight out of twenty-three non-enzymatic glycolysis and PPP-like reactions were six-fold accelerated over their rate in water with this group of amino acids.

### Ribose 5-phosphate forms from 6-phosphogluconate in the presence of cysteine as in the pentose phosphate pathway

Sulfur is an essential component of metabolism, and sulfur-containing amino acids implicated in enzymatic redox catalysis are responsible as well for key structural properties of enzymes. Enzymes of glycolysis, for instance GAPDH, use cysteine in their catalytic center to catalyze reactions that are non-enzymatically catalyzed by Fe(II) (40). Sulfur itself is not directly implicated in the canonical glycolysis and the PPP, but a microbial sulfur-glycolysis has been discovered and may be responsible for major parts of the global sulfur cycle, indicating that the evolution of sulfur metabolism is linked to the evolution of central metabolism (41, 42). Indeed, also the downstream metabolic processes like the Krebs cycle and amino acid biosynthesis depend on sulfur by using FeS clusters in their catalysis (43).

In our screening, sulfur containing amino acids had their most striking effect on the non-enzymatic reactivity of 6-phosphogluconic acid, the central carbohydrate of the PPP. In water, and in the presence of all other amino acids, 6-phosphogluconate was remarkably stable and didn’t yield any substantial product (**Fig. 2A**). In the presence of the sulfur containing amino acids, however, 6-phosphogluconate was converted into other intermediates of the PPP. The metabolite formed at the highest rate was ribose 5-phosphate, the sugar-phosphate that serves as backbone metabolite of RNA and DNA, followed by ribulose 5-phosphate/xylulose 5-phosphate (indistinguishable from each other by the LC-SRM method), and pyruvate (**Fig. 2B**).

We next determined whether cysteine, methionine or both sulfur-containing species were responsible for enhancing 6-phosphogluconate transformations, and found that effects were exclusively explained by cysteine (**Fig. 2C**). As reaction rates in non-enzymatic PPP reactions are strictly pH dependent (31), we tested product formation under different pH regimes in order to set optimal conditions for further characterization of ribose 5-phosphate formation. 50 mM phosphate solutions spanning pH values between 3 and 9 were used. We obtained strongly pH-dependent reaction rate profiles that were distinct for the different products detected, making prominent changes evident and relatively narrow optimal pH values under cysteine conditions, suggestive of catalysis (**Fig. S2**). Other intermediates (erythrose 4-phosphate and 6-phosphogluconolactone) in the PPP were also detected under certain pH conditions (**SI Appendix, Supplementary Information Text**). Maximum quantities of ribose 5-phosphate were observed under mild acidic conditions. Therefore, further experimental conditions were set at pH 5, where cysteine is also stabilized (**Fig. S3**).

Next, we tested alternative sulfur-containing additives to compare their role in the non-enzymatic reaction from 6-phosphogluconate. Ribose 5-phosphate formation rate was differently affected by the oxidation state of the sulfur as well as the other pendant functional groups. While the thiol group (R-SH) was crucial for the activity, the carboxylic and amino groups appeared to play significant roles too as they had important modulatory effects, suggesting a relatively narrow chemical spectrum for drivers of these interconversions, where cysteine effects are moderately exclusive (**Fig. S4**). Oxidized sulfur groups didn’t produce any ribose 5-phosphate (**SI Appendix, Supplementary Information Text**). Finally, we examined the effects of molecular oxygen on this reaction. It was observed that the formation of ribose 5-phosphate did not require a strict anaerobic environment as the reaction occurred too at ambient, aerobic conditions (**Table S2**).

### Cysteine and iron form complementary units of an active bioinorganic complex

At this stage, once the impact of cysteine on 6-phosphogluconate reactivity and ribose 5-phosphate formation was established, we turned to analyze the kinetics in the presence of the metal Fe(II). Can this metal ion account for similar reaction rates as the amino acid promotes? To what extent are these complementary? 800 μM 6-phosphogluconate samples were incubated in 50 mM phosphate solutions containing either FeCl2 or both FeCl2 and cysteine. Already in the case of conditions with the metal ion alone not only the products were the same as those favored by cysteine itself, but the pH-dependence profiles of the reaction rates were similar, aside from slight shifts in the optimum pH values displayed (**Fig. S2**). Remarkably, the maximum formation rate for ribose 5-phosphate still occurred at pH 5. Fe(II) was, comparatively, more efficient than cysteine in promoting ribose 5-phosphate production (**Fig. 3A**).

**Figure 3.**
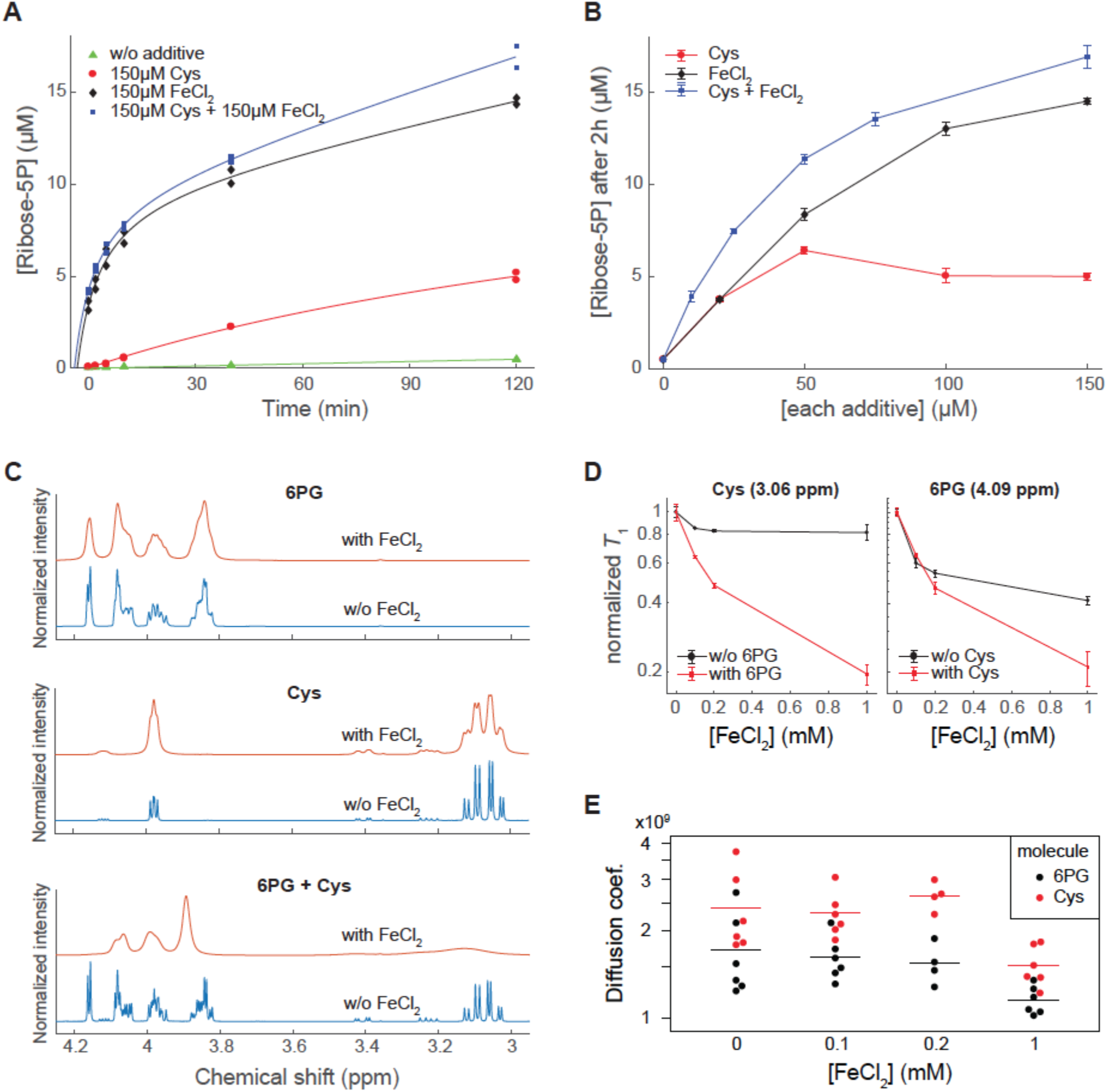
Cysteine and Fe(II) complementary effects on 6PG transformations and the formation of a ternary complex, the ‘minimal enzyme. (A-B) The formation of ribose 5-phosphate at 70°C from 800 μM 6PG in 50 mM phosphate solution pH 5 was monitored for different concentrations of cysteine, different concentrations of Fe(II), and different concentrations of both species combined (molar ratio 1:1). Shown are a comparison of time courses when cysteine and/or Fe(II) were each added at 150 μM (A) and the trends in ribose 5-phosphate concentration at 2h for different individual enhancer concentrations (error bars: mean ± SD) (B). *N* ≥ 3 in all conditions. (C) ^1^H-NMR spectra of 6PG alone, cysteine alone, or the combination of both (20 mM of each analyte) in 50 mM phosphate pH 5/D_2_O solutions, either in the absence or presence of 1 mM FeCl2. The massive disruption and suppression of the ^1^H signal peaks upon addition of Fe(II) to the binary mixture suggests a coordination between cysteine and 6PG in Fe(II) binding. Spectra are shown normalized to the maximum peak intensity per spectrum. (D) T1 relaxation time scaling of 6PG (4.09 ppm) and cysteine (3.06 ppm) signals with increasing Fe(II) concentrations when analyzed separately vs. in combination (conditions as in (C)). (E) Diffusion coefficients measured by diffusion-ordered spectroscopy (DOSY) for cysteine and 6PG molecules (20 mM each) when co-incubated with a mixture of increasing concentrations of Fe(II) and constant, saturating concentrations of phenanthroline, a strong, non-chelating Fe-binding complex. Both molecules showed a noisy but significant decrease in diffusion, a further indication of Fe(II) binding. *N* ≥ 4.

In order to better characterize the underlying dynamics when both chemical species were combined, the analysis was expanded to different concentrations and diverse molar ratios of cysteine to iron. Despite the fact that the activity from cysteine (unlike iron) was shown to quench itself at concentrations above 100 μM, for the whole range of individual concentrations tested the maximum ribose 5-phosphate production was still obtained with the mixture, where the contribution from both components added up (1:1 molar ratio) (**Fig. 3A-B**). Data from different molar ratios of these two species suggests that the combined, net contribution is consistent with a simple model of weighted additive effects rather than a synergy between the amino acid and the metal (**Fig. S5**). Yet, our data indicates that any factor favoring the sole cooccurrence or local proximity between the amino acid and the metal would have sufficed to confer advantage in an early evolutionary stage of primordial biocatalysis.

Since cysteine and iron have a strong binding affinity and are well known to form a variety of complexes that regulate protein function, we investigated if there was any structural indication of a specific combined interaction with 6-phosphogluconate. For this, we made use of ^1^H-NMR spectroscopy, taking advantage of the fact that Fe(II) is paramagnetic and it distorts and suppresses the signal intensities of closely proximal protons from adjacent molecules, in part by reducing the spin-lattice relaxation times of coupled nuclei (*T*_1_) (**Fig. 3C-E; Materials and Methods**). Exposure of 20 mM 6-phosphogluconate to increasing concentrations of Fe(II) in a 50 mM phosphate D_2_O solution at pH 5 confirmed the interaction of the sugar phosphate with the metal ion, as shown previously in unbuffered, aqueous conditions (Keller et al, 2016) (**Fig. 3C**). In particular, the protons bound to C2 and C6 (chemical shifts at 4.09 ppm and 4.16 ppm, respectively) were the most responsive ones (**Fig. 3C**); hence, the interaction was attributed to occur through the carboxylic and phosphate terminal groups of 6-phosphogluconate. Similarly, cysteine demonstrated a fairly strong interaction with Fe(II) at pH 5, as revealed by the smoothing and broadening of its ^1^H-NMR spectral peaks (3.97 ppm and 3.06 ppm), even at relatively low molar fractions of Fe (II) (e.g. with 1 mM Fe(II)) (**Fig. 3C**). In this case, the possible contributions due to interactions through the carboxylic group or the thiol group could not be easily disentangled, given the close spatial proximity between the resonant protons. Strikingly, under the same concentrations as above, Fe(II) dramatically suppressed resonances from both cysteine and 6-phosphogluconate spectra when these species were mixed together (**Fig. 3C**). This indicated the much tighter interaction achieved with the three-components system, further supported by the *T*_1_ relaxation time analyses: shortening of cysteine *T*_1_ relaxation time by Fe (II) became much more pronounced in the presence of 6-phosphogluconate, and vice versa (**Fig. 3D**). In contrast, no significant or much weaker interaction was attained when cysteine was substituted with methionine, or when 6-phosphogluconate was replaced by the analogue glucose 6-phosphate (**Fig. S6**), confirming the specificity of the interaction.

Altogether, the data suggest a cooperative binding between cysteine, Fe(II) and the substrate 6-phosphogluconate, which would form an assembly that could presumably underpin the catalysis. Molecular diffusion measurements carried out by diffusion-ordered spectroscopy (DOSY) ^1^H NMR spectroscopy showed a statistically significant decrease (Mann-Whitney *U* test, α=0.05) in both 6-phosphogluconate and cysteine diffusion coefficients when these were exposed simultaneously to increasing concentrations of Fe(II) in the presence of a relatively bulky, non-chelating Fe(II) ligand such as phenanthroline (**Fig. 3E**). This constituted further evidence to the formation of a (at least transient) reaction-favoring assembly.

### Thermal-dependent reaction specificity of Cys:Fe-driven ribose 5-phosphate formation

So far, the analysis of amino acid-metal influence on the non-enzymatic sugar-phosphate reactivity has been circumscribed to 70°C, a relatively high temperature. Under these conditions, however, temperature is assumed to be a major driver of the net reaction kinetics. Although there are arguments why the origin of metabolism could have happened under thermophilic conditions (44, 45), there are important opposing views. For instance, a high-temperature origin of metabolism would argue that thermophiles could better deal with simple metabolic enzymes, when in fact, they rely on highly structured and advanced enzymes (46). Therefore, we tested if cysteine:Fe(II)-mediated ribose 5-phosphate formation could also proceed at lower temperatures, down to 20°C. 800 μM 6-phosphogluconate was prepared as before in a 50 mM phosphate solution pH 5 containing 75 μM cysteine and 75 μM Fe(II), and the incubation time was extended up to 3 days, depending on temperature, to allow enough time for detectable product formation. Remarkably, cysteine and Fe(II) were able to favor ribose 5-phosphate formation in the whole range of temperatures, even though the reaction rate scaled down as decreasing the temperature, in agreement with the Arrhenius equation (**Fig. 4A-B**). The least-squares fit yielded a value for the activation energy *Ea* of 69 KJ mol^-1^ (±8; 95% confidence bounds). For the same range of temperatures, only very limited formation of ribose 5-phosphate was observed in the absence of the amino acid and the metal, according to a higher energy barrier, estimated to be around 87 KJ mol^-1^ (**Fig. 4A**).

**Figure 4.**
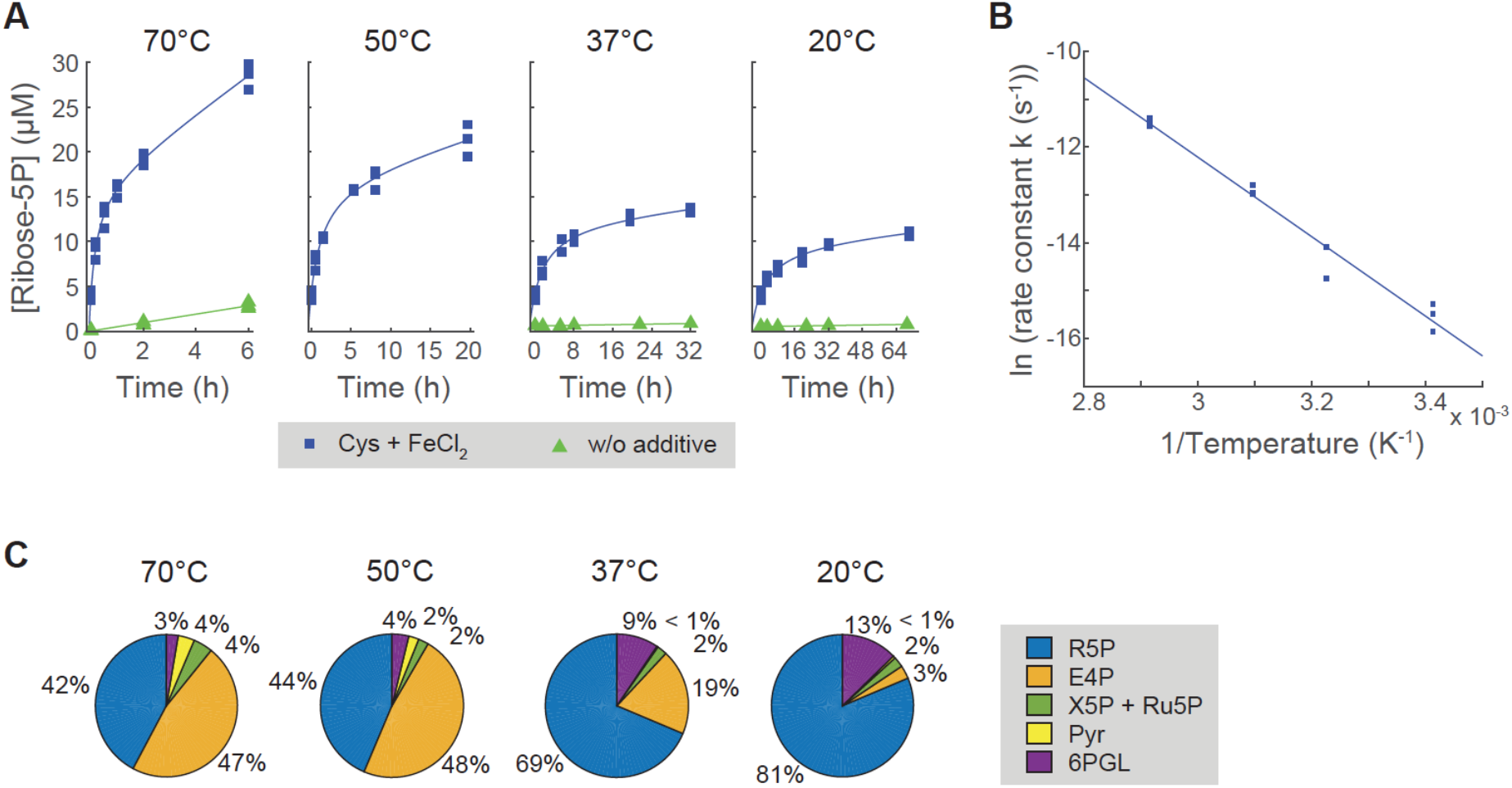
Thermal-dependent non-enzymatic ribose 5-phosphate formation and product specificity from 6PG as in the pentose phosphate pathway. (A) Ribose 5-phosphate concentration over time, formed upon the incubation of 800 μM 6PG in 50 mM phosphate solution pH 5 at different temperatures in the presence of complex-forming 75 μM cysteine and 75 μM FeCl_2_ Notice that time ranges were adapted according to the temperature-scaling kinetics. Experimental data (points; *N* = 3) conform to hyperbolic fits (lines). Trends without the additives are shown for comparison. (B) Initial rate constants calculated from (A) scale exponentially with the absolute temperature, following Arrhenius’ equation. The linear slope in the plot of log-transformed values gives a value for the activation energy Ea of 69 KJ mol^-1^. (C) Long-term product specificity at different temperatures. Relative concentrations of the different detected sugar phosphate products are shown for the last recorded time point per temperature condition from the experiments in (A) (average from *N* = 3 experiments).

Interestingly, temperature-driven changes in reaction rate inversely correlated with product specificity (**Fig. 4C**, **Fig. S7**). While 42% of all 6-phosphogluconate-derived product corresponded to ribose 5-phosphate at relatively late time points under 70°C, this value increased to 81% at 20°C. The relative proportion of other products such as xylulose/ribulose 5-phosphate, pyruvate, and especially erythrose 5-phosphate diminished when decreasing the temperature to ambient temperature. Only 6-phosphogluconolactone was more abundant at lower temperatures, probably due to stability of cyclic structure at lower temperatures thereby trapping the reaction at this intermediate. In consequence, the relative loss of interconversion rate efficiency was compensated by an increased formation of the RNA backbone metabolite Ribose-5-phosphate at low temperatures.

## Discussion

The ‘end-product problem’ describes a classic chicken-egg dilemma for the evolutionary origin of metabolic pathways, which require multiple reaction steps to produce a metabolite. This is a problem for Darwinian selection, as the intermediate steps in a metabolic pathway provide no selective advantage (38). In turn, the “end-product”-forming enzyme can only be selected for after the intermediates are in place. Non-enzymatic, chemical networks might be the basis to overcome this conundrum, by providing an underlying web of subtle preset transformations, upon which Darwinian selection can operate (5). If this hypothesis were true, metabolic pathways that catalyzed a similar set of reactions could come into being multiple times and independently to one another. We noticed that glycolysis and the pentose phosphate pathway (PPP), two central and ancient pathways of central carbohydrate metabolism, are compatible with this hypothesis. First, their topological organization resembles a chemical network that is driven by the most abundant transition metal in Archean sediment, Fe(II) (28, 29, 31, 47). Second, the enzymes that catalyze these pathways in Archaea and Eubacteria have no sequence conservation, and might have come into being independently (23, 39, 48). Despite such a substantial biological evolution, the similarity in the underlying chemistry indicates an ancient origin for these pathways.

Finding prebiotically relevant compounds with catalytic potential on biochemical reactions is one of the big challenges for understanding the origin of metabolism. To date, much of these prebiotically relevant catalytic activities have been ascribed to minerals and metals, while small (bio-)molecules, including amino acids, form only a tiny fraction of it (30, 49–52). Yet, often the prebiotic reactions and pathways described do not map to the structure of extant metabolic networks. In this work we show how single amino acids, and in particular cysteine, can enhance several iron-driven, non-enzymatic reactions between sugar phosphates, resembling extant glycolysis and PPP. Interestingly, cysteine (as well as histidine) has already been implicated in the non-enzymatic catalysis of glycolysis and in particular in acetyl-CoA formation in an earlier study, even though this used huge amounts of compounds and remained mainly qualitative (53). Cysteine was not found in the original Miller’s experiments nor in the Murchison meteorite and has for long been criticized as a prebiotically relevant amino acid. However, more recent reanalysis of prebiotic synthesis experiments under simulated prebiotic atmosphere including H2S, report abundant sulfur-containing amino acids (54); see also (55). While one might indeed debate the relevance of Miller’s experiments for the evolution of early enzymes, there are however important biological arguments that support the early importance of cysteine: All species universally use cysteine in their essential metabolic enzymes, indicating that it was available when the metabolic network evolved. Cysteine hence was prevalent in biological evolution, and additionally, other sulphur containing molecules could have fulfilled the role (56).

The role of cysteine could be that of a nucleophile as observed in the case of the enzyme aldehyde dehydrogenase (57). Although the exact chemistry behind the non-oxidative reactions remains elusive, the described chemical system establishes the intermediates of the PPP and Embden-Meyerhof pathways via a series of reactions observed in Maillard reactions as well as thiol-mediated oxidation reactions (58). Cysteine sulfur condenses with a carbonyl functionality allowing for the facile dehydrogenation (**Fig 5**). The resulting thioester undergoes hydrolysis to give the corresponding carboxylic acid.

**Figure 5.**
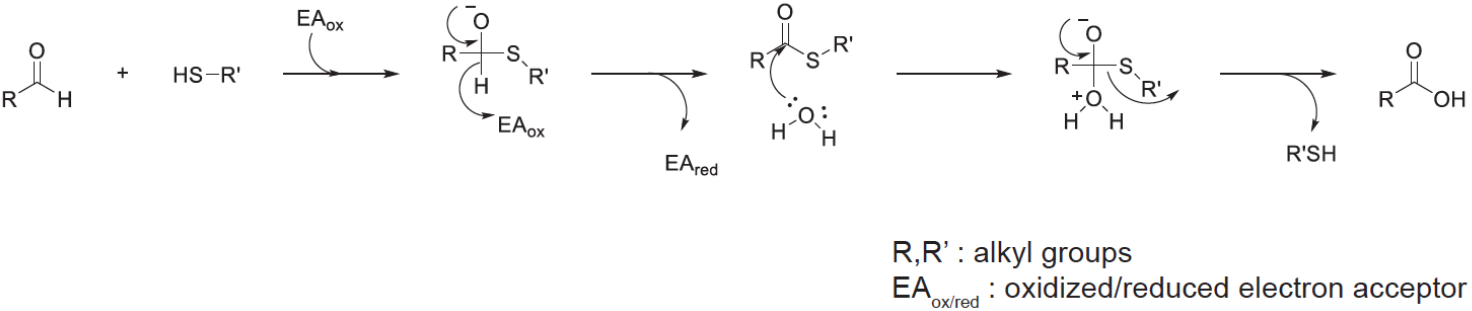
Proposed mechanism for thiol catalyzed oxidation of aldose sugar to sugar acid. EA could be an electron acceptor such as oxygen or Fe(III) generated *in situ*.

In addition to this observed organocatalysis by cysteine, we also observe a complex formation between different components - a sugar-phosphate, a metal and cysteine. The amino acid ligand, in addition to acting as a surrogate of metal catalysis, might also act as a scaffold between the substrate and the metal. This could be regarded as a possibility for how a transition from simple metal-ligand catalyst to longer peptides based on metal centers could have taken place (49, 59).

The prevalence of life across a wide range of temperatures shows the robustness of its cellular components to high thermal energies. This resistance to decomposition may have acted as a major initial selection criterion along with catalytic rates (60). Pre-existent chemical scaffolds would have interacted with the already present metal catalysts and those (sub)networks contributing with intermediates that reinforced metal catalysis or reaction stability were further selected and retained in nascent metabolic pathways. Single amino acids (or small peptides) could have embodied those attributes enucleating on metal centers, at the time that they could have later relegated metals making metabolism less dependent on environmental catalysts (61, 62). By being a rate-limiting reaction, even a slight enhancement of ribose 5-phosphate production from 6-phosphogluconate under a diversity of conditions with moderate specificity, could have been an excellent asset in mild microenvironments during the early evolution of metabolism. At that era, time was presumably not as limiting as it was the ability to consolidate autonomous procurement of specific metabolites such as ribose-5-phosphate under conditions where other metabolites of the metabolic network, as well as early components that led to the evolution of the transcription and translational machinery, could persist too. From this perspective, it becomes plausible that early enzymes evolved stepwise and did not need to provide high specificity or yield to make a positive impact on an early metabolism.

In summary, we have shown that single-amino acids can alter a broad range of activity within a non-enzymatic, metal promoted network that resembles glycolysis and the pentose phosphate pathway. Cysteine, as sulfur containing proteinogenic amino acid in particular formed a ternary complex with Fe(II) and 6-phosphogluconate, and acting as a ‘minimal enzyme’, promoted the formation of ribose 5-phosphate, in analogy to a rate limiting reaction of the oxidative PPP. Moreover, we noted that mundane (20°C) temperatures substantially increased the specificity of ribose 5-phosphate formation. Our results hence show that reactions of metal-catalyzed networks can be replaced with simple amino acid-based catalysis, and hence propose a scenario in which non-enzymatic, fuzzy chemical networks serve as a template for the selection of enzymatic catalysts in the early evolution of metabolism. This scenario implies a solution for the ‘end product problem’, because in the presence of a pre-existing metal driven reaction network, not all enzymes that form a metabolic pathway need to come into being at once to enable the formation of a selectable product. Further, this model could explain why metabolic network topologies remain conserved upon the evolution of enzymatic catalysis.

## Materials and Methods

### Materials

All chemicals were obtained from Sigma-Aldrich. UPLC/MS grade acetonitrile (AcCN) and water were purchased from Greyhound Chemicals. (see **SI Appendix, Supplementary Methods** for further details)

### Sample preparation for reaction kinetics experiments

Samples consisting of individual sugar phosphates dissolved (in the μM range) in 200 μL aqueous solutions (either in 50 mM phosphate or UPLC/MS grade water) were prepared fresh. These were placed in 2ml screw thread amber autosampler glass vials (Agilent Technologies) and sealed in an anaerobic chamber (Coy Laboratory Products) following 3 vacuum/N2 injection cycles to remove oxygen content. Samples were then incubated in a water bath at the indicated temperature for different times, after which reactions were stopped by immediate transference to ice. Samples were stored at −20°C until thawed for LC-MS/MS measurements. The incubation time points were specifically set for each metabolite based on its previously reported stability (29, 31); these were typically defined at increasing time intervals and spanned for up to 6h in the experiments at 70°C.

### LC-MS/MS and quantification of sugar phosphate interconversions

Samples were analyzed for sugar-phosphate content by a tandem liquid chromatography-mass spectroscopy (LC-MS/MS) system, using a triple-quadrupole device (Agilent 6460) operating in SRM mode (parameter specifications and metabolite ion transitions defined as in (29); see also (63). A reverse-phase, C8 chromatographic column (Zorbax SB-C8 RRHD 2.1 × 100 mm 1.8 μm from Agilent) was used for the separation, with two binary acetonitrile/water mobile phases (Buffer A: 90% Water, 10% AcCN; Buffer B: 50% Water, 50% AcCN) applied in gradient, both containing 750 mg/l octylammonium acetate as ion-pairing reagent. Details were as followed: 7.5-min cycle at isocratic 0.6 ml/min flow, with elution with 5% Buffer B for 3.5 min, followed by a 2.5-min ramp to 70% of B, 0.5 min at 80% B and 1-min re-equilibration to 5% B. Individual sugar phosphates could be resolved by retention time as well as fragmentation pattern, as contrasted with external standards. Only Ru5P and X5P (229 -> 97 m/z) and 2PG and 3PG (185 -> 97 m/z) could not be discriminated, and were considered together in the analysis. To avoid possible biases in quantification due to long-batch effects (e.g. variations in instrument sensitivity), sample triplicates were randomized in the autosampler positions before MS measurements.

MS/MS data were analyzed with the quantitative MassHunter software (Agilent), with manual supervision of the automated signal-peak integration. Several replicates of sugar-phosphate external-standard dilution series allowed to translate signal intensities into absolute concentrations. This was achieved with less accuracy in the case of E4P due to its broad, low-intensity peak, as described in previous work (64), and because of its remarkably poor purity (50%) in the commercially available standard. Also, the lack of a commercial standard for 6PGL forced us to estimate its concentration from the rapid favored equilibrium with 6PG occurring at a strong acidic pH. Concentration time courses were reconstructed with MATLAB. Only cases with over-background product detection showing a congruent increase between triplicates were considered as significant transformations and analyzed for the kinetics. Reaction kinetics were fitted according to one of the following models, depending on the adequacy to the experimental data (R^2^ values): a least-squares linear production model, a saturating hyperbolic model or sigmoidal Gompertz function, or a mixed model comprising a first linear phase of maximum (initial) growth rate and a slow exponential decay. Rates, when reported, are shown in terms of the maximum (initial) slopes of the reaction kinetics fits.

### NMR studies of metabolite-iron interaction

Samples containing a final metabolite concentration of 20 mM were prepared in deuterated water (D_2_O) containing 0.05 mM trimethylsilyl propanoic acid (TSP) as internal standard, sodium azide and 50 mM sodium phosphate at a pH 5. FeCl2 was added at concentrations of 0, 0.1, 0.2 and 1 mM. NMR experiments were carried out using an AVANCE II+ (Bruker, Rheinstetten, Germany) NMR spectrometer operating at 500.13 MHz for the ^1^H frequency using a 5 mm TXI probe. Basic 1D spectra were collected using a solvent suppression pulse sequence based on a onedimensional version of the nuclear Overhauser effect spectroscopy (NOESY) pulse sequence to saturate the residual ^1^H water signal (relaxation delay = 2 s, t1 increment = 3 μs, mixing time = 150 ms, solvent presaturation applied during the relaxation time and the mixing time). 128 transients were collected into 16 K data points over a spectral width of 12 ppm at 320 K.

Longitudinal relaxation times (*T*_1_) were measured using the Inversion recovery pulse sequence, 180-τ-90, with pulse spacing τ having the following values: 0.01 s, 0.02 s, 0.03 s, 0.05 s, 0.1 s, 0.25 s, 0.5 s, 0.75 s, 1 s, 1.5 s, 2 s, 4 s, 8 s, 15 s, 20 s, 25 s. The pulse repetition time was set at 20 s. Eight transients were collected into 16 K data points over a spectral width of 20 ppm at 320 K.

Diffusion coefficients were measured using a bipolar gradient pulse pair with a spoil gradient pulse (stebpgp1s), with 16 incremental steps in the gradient strength linearly ramped from 2% to 95% of the maximum gradient strength. 16 Scans per increment step were collected into 16K data points over a spectra width of 19 ppm at 320 K. The gradient pulse length was set at 1000 μs, while the time between pulses at 0.1 sec. The pulse repetition time was set at 2 s.

1D NMR spectra were processed using TopSpin v.3.2 (Bruker, Rheinstetten, Germany). Free induction decays were Fourier transformed following multiplication by a line broadening of 1 Hz, and referenced to TSP at 0.0 ppm. Spectra were automatically phase and baseline corrected. For the relaxation times and diffusion coefficients, data were analyzed using the T1/T2 Relaxation module present in TopSpin. All experiments were carried out at least by triplicate.

## Acknowledgements

We thank members of Ralser’s group for valuable discussions and critical comments, Olivier Languin-Cattoën and Maria Ouvarova for help in the non-enzymatic reaction screening, and Dr. Jae-Hun Jeoung (Humboldt University Berlin) for assistance in carrying out anaerobic reactions. This work was supported by the Francis Crick Institute which receives its core funding from Cancer Research UK (FC001134), the UK Medical Research Council (FC001134), and the Wellcome Trust (FC001134), and received specific support from the Wellcome Trust (200829/Z/16/Z) and the Ministry of Education and Research (BMBF), as part of the National Research Node ‘Mass spectrometry in Systems Medicine (MSCoresys), under grant agreement 031L0220A. Work in the JLG lab is supported by the Medical Research Council UK (MR/P011705/1, MC_UP_A090_1006). GP acknowledges support by a Talento program fellowship from Comunidad de Madrid.

## References

1. G. Wächtershäuser, Evolution of the first metabolic cycles. Proc. Natl. Acad. Sci. U. S. A. 87, 200–204 (1990).

2. A. Lazcano, S. L. Miller, On the origin of metabolic pathways. J. Mol. Evol. 49, 424–431 (1999).

3. H. J. Morowitz, J. D. Kostelnik, J. Yang, G. D. Cody, The origin of intermediary metabolism. Proc. Natl. Acad. Sci. U. S. A. 97, 7704–7708 (2000).

4. L. E. Orgel, The implausibility of metabolic cycles on the prebiotic Earth. PLoS Biol. 6, e18 (2008).

5. M. Ralser, The RNA world and the origin of metabolic enzymes. Biochem. Soc. Trans. 42, 985–988 (2014).

6. J. C. Xavier, W. Hordijk, S. Kauffman, M. Steel, W. F. Martin, Autocatalytic chemical networks at the origin of metabolism. Proc. Biol. Sci. 287, 20192377 (2020).

7. N. Kitadai, S. Maruyama, Origins of building blocks of life: A review. Geoscience Frontiers 9, 1117–1153 (2018).

8. S. L. McKnight, On Getting There from Here. Science 330, 1338–1339 (2010).

9. R. J. DeBerardinis, C. B. Thompson, Cellular metabolism and disease: what do metabolic outliers teach us? Cell 148, 1132–1144 (2012).

10. J. P. DeLong, J. G. Okie, M. E. Moses, R. M. Sibly, J. H. Brown, Shifts in metabolic scaling, production, and efficiency across major evolutionary transitions of life. Proc. Natl. Acad. Sci. U. S. A. 107, 12941–12945 (2010).

11. R. A. Notebaart, et al., Network-level architecture and the evolutionary potential of underground metabolism. Proc. Natl. Acad. Sci. U. S. A. 111, 11762–11767 (2014).

12. S. Varahan, A. Walvekar, V. Sinha, S. Krishna, S. Laxman, Metabolic constraints drive self-organization of specialized cell groups. Elife 8 (2019).

13. M. Lempp, et al., Systematic identification of metabolites controlling gene expression in E. coli. Nat. Commun. 10, 4463 (2019).

14. V. P. Skulachev, The laws of cell energetics. Eur. J. Biochem. 208, 203–209 (1992).

15. G. Piedrafita, M. A. Keller, M. Ralser, The Impact of Non-Enzymatic Reactions and Enzyme Promiscuity on Cellular Metabolism during (Oxidative) Stress Conditions. Biomolecules 5, 2101–2122 (2015).

16. E. Smith, H. J. Morowitz, Universality in intermediary metabolism. Proc. Natl. Acad. Sci. U. S. A. 101, 13168–13173 (2004).

17. J. Peretó, Out of fuzzy chemistry: from prebiotic chemistry to metabolic networks. Chem. Soc. Rev. 41, 5394–5403 (2012).

18. M. Ralser, An appeal to magic? The discovery of a non-enzymatic metabolism and its role in the origins of life. Biochem. J 475, 2577–2592 (2018).

19. L. E. Orgel, Self-organizing biochemical cycles. Proc. Natl. Acad. Sci. U. S. A. 97, 12503–12507 (2000).

20. R. Fani, The Origin and Evolution of Metabolic Pathways: Why and How did Primordial Cells Construct Metabolic Routes? Evolution: Education and Outreach 5, 367–381 (2012).

21. R. Tanaka, M. Csete, J. Doyle, Highly optimised global organisation of metabolic networks. Syst. Biol. 152, 179–184 (2005).

22. H. Jeong, B. Tombor, R. Albert, Z. N. Oltvai, A. L. Barabási, The large-scale organization of metabolic networks. Nature 407, 651–654 (2000).

23. A. H. Romano, T. Conway, Evolution of carbohydrate metabolic pathways. Res. Microbiol. 147, 448–455 (1996).

24. T. Geisberger, et al., Evolutionary Steps in the Analytics of Primordial Metabolic Evolution. Life 9 (2019).

25. M. A. Keller, P. C. Driscoll, C. B. Messner, M. Ralser, 1H-NMR as implemented in several origin of life studies artificially implies the absence of metabolism-like non-enzymatic reactions by being signal-suppressed. Wellcome Open Res 2, 52 (2018).

26. P. Van Cappellen, E. D. Ingall, Redox Stabilization of the Atmosphere and Oceans by Phosphorus-Limited Marine Productivity. Science 271, 493–496 (1996).

27. J. C. Walker, P. Brimblecombe, Iron and sulfur in the pre-biologic ocean. Precambrian Res. 28, 205–222 (1985).

28. O. J. Rouxel, A. Bekker, K. J. Edwards, Iron isotope constraints on the Archean and Paleoproterozoic ocean redox state. Science 307, 1088–1091 (2005).

29. M. A. Keller, A. V. Turchyn, M. Ralser, Non-enzymatic glycolysis and pentose phosphate pathway-like reactions in a plausible Archean ocean. Mol. Syst. Biol. 10 (2014).

30. K. B. Muchowska, et al., Metals promote sequences of the reverse Krebs cycle. Nature Ecology & Evolution 1, 1716–1721 (2017).

31. M. A. Keller, et al., Conditional iron and pH-dependent activity of a non-enzymatic glycolysis and pentose phosphate pathway. Sci Adv 2, e1501235 (2016).

32. S. J. Varma, K. B. Muchowska, P. Chatelain, J. Moran, Native iron reduces CO2 to intermediates and end-products of the acetyl-CoA pathway. Nat Ecol Evol 2, 1019–1024 (2018).

33. K. B. Muchowska, S. J. Varma, J. Moran, Synthesis and breakdown of universal metabolic precursors promoted by iron. Nature 569, 104–107 (2019).

34. M. Preiner, et al., A hydrogen-dependent geochemical analogue of primordial carbon and energy metabolism. Nat Ecol Evol (2020) https:/doi.org/10.1038/s41559-020-1125-6.

35. C. B. Messner, P. C. Driscoll, G. Piedrafita, M. F. L. De Volder, M. Ralser, Nonenzymatic gluconeogenesis-like formation of fructose 1,6-bisphosphate in ice. Proceedings of the National Academy of Sciences 114, 7403–7407 (2017).

36. R. S. Ronimus, H. W. Morgan, Distribution and phylogenies of enzymes of the Embden-Meyerhof-Parnas pathway from archaea and hyperthermophilic bacteria support a gluconeogenic origin of metabolism. Archaea 1, 199–221 (2003).

37. A. J. Coggins, M. W. Powner, Prebiotic synthesis of phosphoenol pyruvate by α-phosphorylation-controlled triose glycolysis. Nat. Chem. (2016) https:/doi.org/10.1038/nchem.2624 (November 29, 2016).

38. N. H. Horowitz, On the Evolution of Biochemical Syntheses. Proc. Natl. Acad. Sci. U. S. A. 31, 153–157 (1945).

39. S. M. Kengen, A. J. M. Stams, W. M. de Vos, Sugar metabolism of hyperthermophiles. FEMS Microbiol. Rev. 18, 119–137 (1996).

40. S. V. Antonyuk, R. R. Eady, R. W. Strange, The structure of glyceraldehyde 3-phosphate dehydrogenase from Alcaligenes xylosoxidans at 1.7 Å resolution. Section D: Biological… (2003).

41. K. Denger, et al., Sulphoglycolysis in Escherichia coli K-12 closes a gap in the biogeochemical sulphur cycle. Nature 507, 114–117 (2014).

42. A. K. Felux, D. Spiteller, Entner–Doudoroff pathway for sulfoquinovose degradation in Pseudomonas putida SQ1. Proceedings of the (2015).

43. S. J. Lloyd, H. Lauble, G. S. Prasad, C. D. Stout, The mechanism of aconitase: 1.8 A resolution crystal structure of the S642a:citrate complex. Protein Sci. 8, 2655–2662 (1999).

44. M. C. Weiss, et al., The physiology and habitat of the last universal common ancestor. Nat Microbiol 1, 16116 (2016).

45. J. Wiegel, A. W. W. Michael, Thermophiles: The Keys to the Molecular Evolution and the Origin of Life? (CRC Press, 1998).

46. G. Gianese, F. Bossa, S. Pascarella, Comparative structural analysis of psychrophilic and meso-and thermophilic enzymes. Proteins: Struct. Funct. Bioinf. 47, 236–249 (2002).

47. M. A. Saito, D. M. Sigman, F. M. M. Morel, The bioinorganic chemistry of the ancient ocean: the co-evolution of cyanobacterial metal requirements and biogeochemical cycles at the Archean–Proterozoic boundary? Inorganica Chim. Acta 356, 308–318 (2003).

48. A. Stincone, et al., The return of metabolism: biochemistry and physiology of the pentose phosphate pathway. Biol. Rev. Camb. Philos. Soc. 90, 927–963 (2015).

49. L. Belmonte, S. S. Mansy, Metal Catalysts and the Origin of Life. Elements 12, 413–418 (2016).

50. R. M. Hazen, D. A. Sverjensky, Mineral surfaces, geochemical complexities, and the origins of life. Cold Spring Harb. Perspect. Biol. 2, a002162 (2010).

51. C. E. Cornell, et al., Prebiotic amino acids bind to and stabilize prebiotic fatty acid membranes. Proc. Natl. Acad. Sci. U. S. A. 116, 17239–17244 (2019).

52. C. Huber, F. Kraus, M. Hanzlik, W. Eisenreich, G. Wächtershäuser, Elements of metabolic evolution. Chemistry 18, 2063–2080 (2012).

53. M. Shimizu, A. Yamagishi, K. Kinoshita, Y. Shida, T. Oshima, Prebiotic origin of glycolytic metabolism: histidine and cysteine can produce acetyl CoA from glucose via reactions homologous to non-phosphorylated Entner-Doudoroff pathway. J. Biochem. 144, 383–388 (2008).

54. E. T. Parker, et al., Prebiotic synthesis of methionine and other sulfur-containing organic compounds on the primitive Earth: a contemporary reassessment based on an unpublished 1958 Stanley Miller experiment. Origins of Life and Evolution of Biospheres 41, 201–212 (2011).

55. B. N. Khare, C. Sagan, Synthesis of cystine in simulated primitive conditions. Nature 232, 577–579 (1971).

56. A. L. Weber, S. L. Miller, Reasons for the occurrence of the twenty coded protein amino acids. J. Mol. Evol. 17, 273–284 (1981).

57. Z. J. Liu, et al., The first structure of an aldehyde dehydrogenase reveals novel interactions between NAD and the Rossmann fold. Nat. Struct. Biol. 4, 317–326 (1997).

58. H. E. Nursten, The Maillard Reaction: Chemistry, Biochemistry and Implications (Royal Society of Chemistry, 2007).

59. L. Belmonte, et al., Cysteine containing dipeptides show a metal specificity that matches the composition of seawater. Phys. Chem. Chem. Phys. 18, 20104–20108 (2016).

60. G. Piedrafita, F. Montero, F. Morán, M. L. Cárdenas, A. Cornish-Bowden, A simple self-maintaining metabolic system: robustness, autocatalysis, bistability. PLoS Comput. Biol. 6 (2010).

61. K. Ruiz-Mirazo, A. Moreno, Basic autonomy as a fundamental step in the synthesis of life. Artif. Life 10, 235–259 (2004).

62. G. Piedrafita, P.-A. Monnard, F. Mavelli, K. Ruiz-Mirazo, Permeability-driven selection in a semi-empirical protocell model: the roots of prebiotic systems evolution. Sci. Rep. 7, 3141 (2017).

63. M. M. C. Wamelink, et al., Quantification of sugar phosphate intermediates of the pentose phosphate pathway by LC–MS/MS: application to two new inherited defects of metabolism. Journal of Chromatography B 823, 18–25 (2005).

64. B. Luo, K. Groenke, R. Takors, C. Wandrey, M. Oldiges, Simultaneous determination of multiple intracellular metabolites in glycolysis, pentose phosphate pathway and tricarboxylic acid cycle by liquid chromatography–mass spectrometry. Journal of Chromatography A 1147, 153–164 (2007).

