## Supplementary Information for "Single amino acid-promoted reactions link a non-enzymatic chemical network to the early evolution of enzymatic pentose phosphate pathway"

**This PDF file includes:**

Supplementary Methods

Supplementary Text

Figures S1 to S7

Tables S2 to S10

SI References

**Other supplementary materials for this manuscript include the following:**

Table S1

### Supplementary Methods

*Materials.* All chemicals were obtained from Sigma-Aldrich. ULC/MS grade acetonitrile (AcCN) and water were purchased from Greyhound Chemicals. The anaerobic experiments from the oxygen effect studies were carried out in a glove box (model B; COY Laboratory Products) under an atmosphere of 95% $N_2$/5% $H_2$(Air-Liquide GmbH). All the solutions used were degassed in a stoppered bottle/tube with a screw cap with butyl rubber septum by successive cycles of evacuating and $N_2$ gas flushing at a Schlenk line. The content of $^{56}Fe$ was measured using an Agilent 7900 ICP-MS instrument fitted with Ni cones and a MicroMistnebulizer in $H_2$ mode.

*Cysteine quantitation*. Cysteine was prepared at 400 μM in 50 mM phosphate solutions at the indicated pHs, and incubated at 70°C for different times under low-oxygen conditions, following the protocol described above for sugar phosphates. Sample vials were cooled by transference to ice (but avoiding freezing) and thereafter cysteine levels quantified by colorimetric assay using Ellman's reagent (1), a free sulfhydryl-reactive chemical yielding a measurable yellow product: A stock solution of 5,5'-dithiobis-(2-nitrobenzoic acid) (DTNB) was added to the samples, to a final concentration of 70 μg/mL. Upon short vortexing and 10-min incubation at room temperature, samples were transferred into 96-well plates and the absorbance at 412 nm measured spectrophotometrically (Infinite 200 PRO microplate reader from Tecan). A standard dilution series made from freshly prepared solutions of known cysteine concentrations was used to obtain absolute concentrations.

### Supplementary Information Text

*pH-dependent product formation of cysteine-driven 6-phosphogluconate reactions:*

Since different products apart from ribose 5-phosphate were accessible from 6-phosphogluconate in (unbuffered) aqueous conditions at 70°C, we inquired how pH changes could affect the reaction specificity and whether the pH would allow tuning the rate efficiency and cysteine preference to form one product over other. Here we comment on the optimal pH for reactivity, and address whether the pH optimum is reaction or product-specific (Fig. S2). Samples containing 800 μM 6-phosphogluconate and 400 μM cysteine were prepared in 50 mM phosphate solutions spanning a wide range of pH values (between 3 and 9). Product yields after 6 h incubation at 70°C turned out to be very dependent on pH (Fig. S2). While pyruvate was preferentially formed at alkaline pHs, intermediate or mild acidic conditions (around pH 5) were the most favorable for the formation of higher-order sugar species from the PPP, i.e. 5-carbon sugar phosphates, and predominantly ribose 5-phosphate. Additionally, other PPP intermediates not found in the original unbuffered aqueous conditions were detected at mild and strong acidic pHs: erythrose 4-phosphate and 6-phosphogluconolactone, respectively.

Altogether, although product-specific, pH-dependent reaction rate profiles with cysteine were mostly different than those obtained in control conditions (without cysteine) and resulted much more prominent, with relatively narrow optimal pH values, suggestive of catalysis (Fig. S2). Only in the case of 6-phosphogluconolactone, production yields were independent of cysteine (Fig. S2)

– indeed, the formation of this species at pH 3 is consistent with favored proton-driven dehydration of 6-phosphogluconate under acidic conditions. Now, the distinguishable modes of response to pH obtained for the different products may point to alternative driven reactions pathways, but could also result from multi-step sequential chemistry. In this context, we chose to focus the analysis on ribose 5-phosphate (and the other -- less abundant -- pentose phosphates) due to their particular relevance for being the metabolites most immediately related to 6-phosphogluconate (at least in vivo), and thus more likely as primary products of catalysis, as for showing an optimum formation rate around pH 5, far from alkaline conditions where cysteine was found more reactive (Fig. S3).

*Role of cysteine functional groups in product formation rate:*

In order to study the role of cysteine functional groups in the sugar conversion performance we design a time-course experiment with a battery of cysteine analogues (Fig. S4). 6-phosphogluconate-derived ribose 5-phosphate formation was analyzed in the presence of each of the following molecular analogues of cysteine (again, at a concentration of 400 μM): Isomers or close homologues bearing the same functional groups (D-cysteine, DL-homocysteine and reduced glutathione (GSH)), structural analogues differing in one or several functional groups (L-serine, cysteamine, β-mercaptoethanol and 3-mercaptopropionic acid), and oxidized cysteine derivatives (cystine, oxidized glutathione (GSSG) and cysteine sulfonic acid) were tested. The results demonstrated that the thiol group (R-SH) was crucial for cysteine activity (Fig. S4). No significant formation of ribose 5-phosphate was detected with the alcohol analogue L-serine, nor with the disulfide-based compounds after 6h incubation with 800 μM 6-phosphogluconate. Conversely, irrespective of slight structural differences, all the thiol-containing analogues mimicked cysteine effects. Interestingly, however, the presence of the carboxylic group was a second feature of advantage, with cysteamine and β-mercaptoethanol showing only modest enhancement effects (Fig. 2C). In addition, the enhanced reaction rate with 3-mercaptopropionic acid would suggest that the local environment of the carboxylic group is also important. These results point to a relatively specific and complex set of conditions required for catalysis even in a small molecule such as an amino acid.

**Supplementary figures and tables**

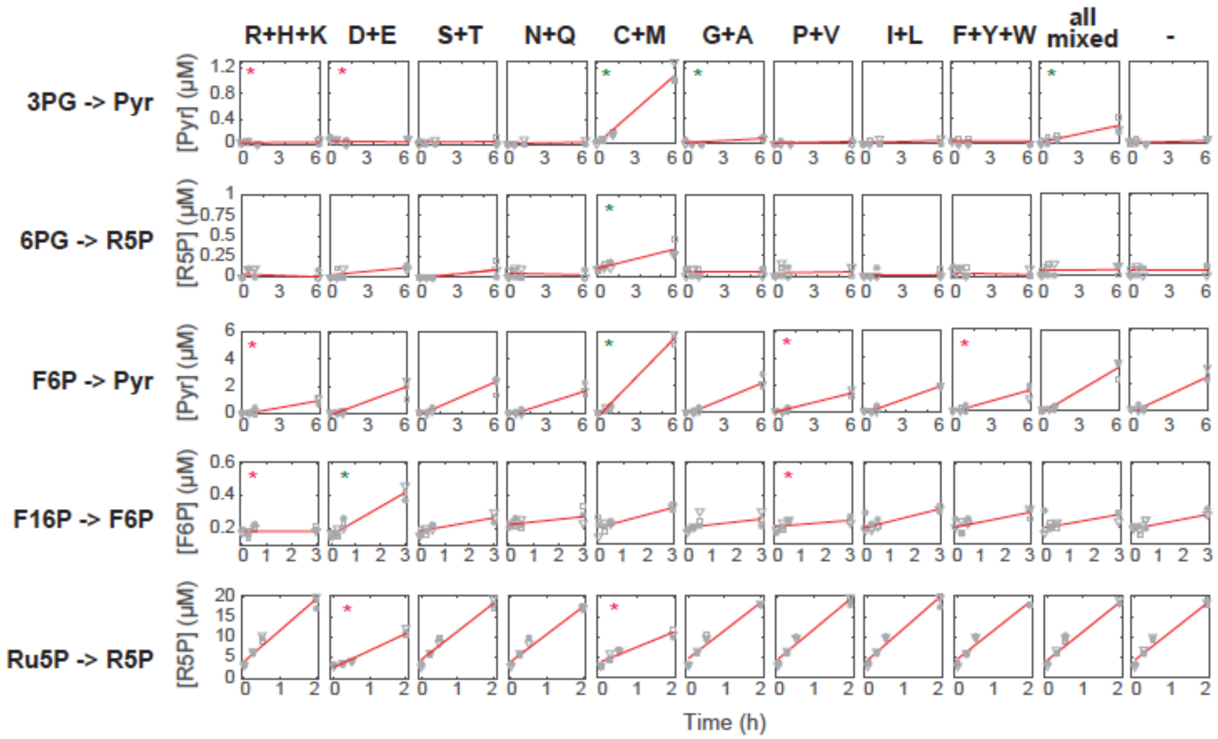


**Fig. S1. Representative examples of non-enzymatic sugar interconversions modulated by amino acids.** Product formation time courses obtained at 70°C from 100 µM substrate in the presence of different subgroups of amino acids (at a total concentration of 400 µM) are shown for different prototypic reactions: diverse transformations accelerated by the sulfur-containing amino acids cysteine and methionine, some of which are negatively affected by at least another group of amino acids (top three panel rows); F1,6BP dephosphorylation, enhanced by negatively-charged amino acids, with antagonistic effects from positively-charged amino acids (fourth set of panels); a pentose-phosphate isomerization, where the role of cysteine and methionine is actually as inhibitors (bottom panel row). Asterisks indicate reaction rates that were significantly different – either higher (green) or lower (red) – than in the aqueous control without amino acids (last column of panels) (Wilcoxon rank-sum test, $p$=0.1). “all mixed” stands for samples containing all 20 amino acids at a total concentration of 400 µM (i.e. 20 µM each). Data points shown in grey ($N$ = 3 independent experiments).

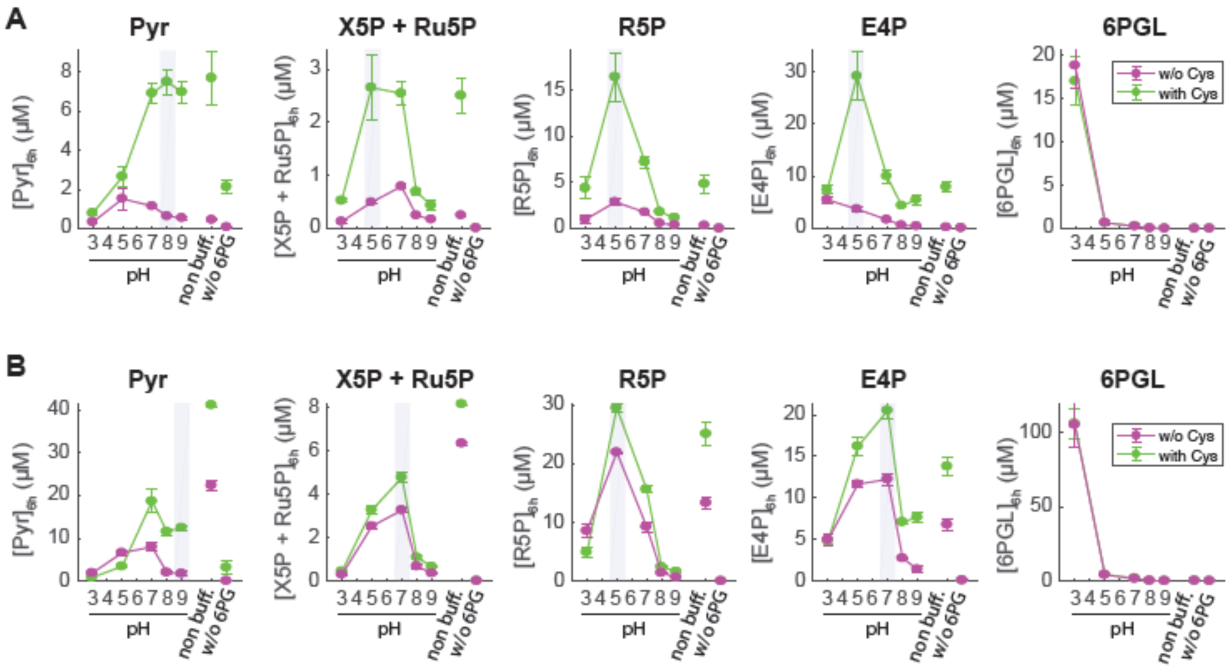


**Fig. S2. pH-dependent activity of cysteine on the formation of different products from 6-phosphogluconate (6PG).** (A) 100 µM 6PG was incubated at 70°C in 50 mM phosphate solution at different pHs, either in the absence (purple lines) or presence (green lines) of 400 µM cysteine. The concentrations of the sugar phosphate products detected and quantified by LC/MS after 6h incubation are displayed. The optimum pH range where the reaction enhancement by cysteine was larger is highlighted as a shaded, grey region where applicable. A control consisting of 100 µM 6PG in unbuffered (aqueous) conditions, as well as a negative control without 6PG are shown for comparison. (B) Same protocol was followed but in the presence of a metal ion, 200 µM $FeCl_2$, either without (purple lines) or with 400 µM cysteine co-present (green lines). Data shown as mean ± SD ($N \geq 3$ in all conditions).

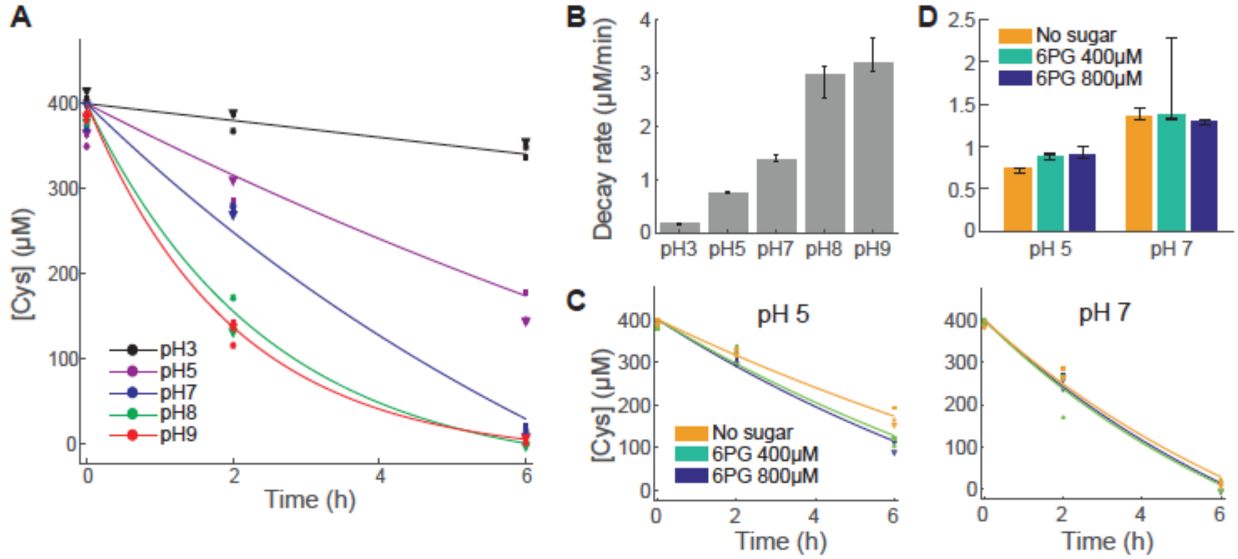


**Fig. S3. Stability of cysteine at 70°C under different experimental conditions.** (A) Time evolution of cysteine concentration during incubation in 50 mM phosphate solution at different pHs. Lines correspond with exponential decay fits. Corresponding rates are shown in (B): mean ± SD ($N \geq 3$). (C) For those conditions of intermediate pH, cysteine time courses were reanalyzed both in absence and presence of increasing concentrations of the substrate, 6PG, with just very slight changes detected. Corresponding rates are shown in (D): mean ± SD ($N \geq 3$). In all experiments cysteine concentration was quantified spectrophotometrically using Ellman's reagent (see Materials and Methods).

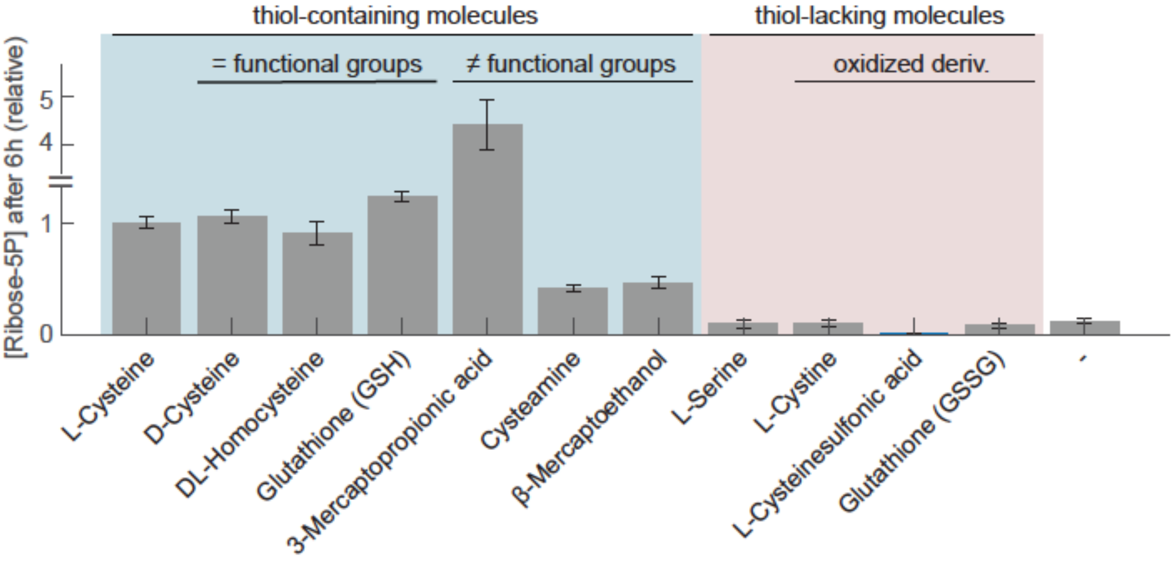


**Fig. S4. Ribose 5-phosphate yields obtained from 6PG with different cysteine analogues.** 800 µM 6PG was incubated at 70°C in a 50 mM phosphate solution pH 5 containing 400 µM of cysteine analogue. The concentration of R5P formed after 6h is shown - relative to the 400 µM cysteine condition - for isomers or close homologues bearing the same functional groups (i.e. D-cysteine, DL-homocysteine and reduced glutathione (GSH)), structural analogues differing in one or several functional groups (3-mercaptopropionic acid, cysteamine, β-mercaptoethanol and L-serine), and oxidized cysteine derivatives (L-cystine, cysteine sulfonic acid and oxidized glutathione (GSSG)), evidencing the importance of the thiol group. Error bars represent mean ± SD ($N = 3$).

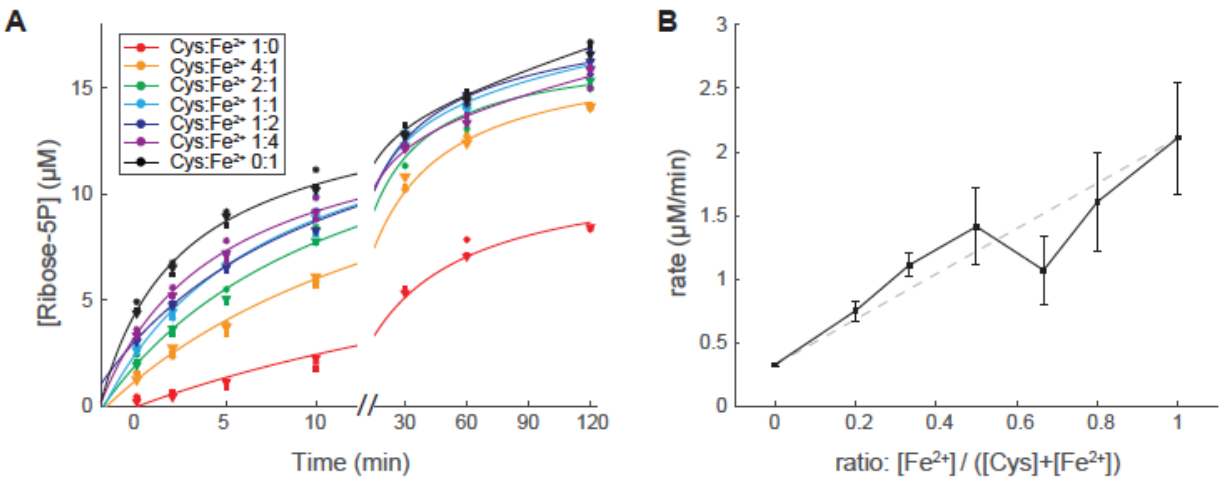


**Fig. S5. Ribose 5-phosphate formation rates with different molar ratios of amino acid and metal.** 800 µM 6PG was incubated at 70°C in 50 mM phosphate solution pH 5 containing different molar ratios of cysteine and $FeCl_2$ but at same total concentration, 150 µM. (A) Detailed time courses in R5P formation are shown. Lines represent best hyperbolic fits. $N = 3$ per condition. (B) Estimated initial rates (error bars: mean ± SD) are plotted as a function of the ratio between both additives. The grey dashed line represents the null-model expectation of weighted additive contributions with no patent inter-species dynamic interaction.

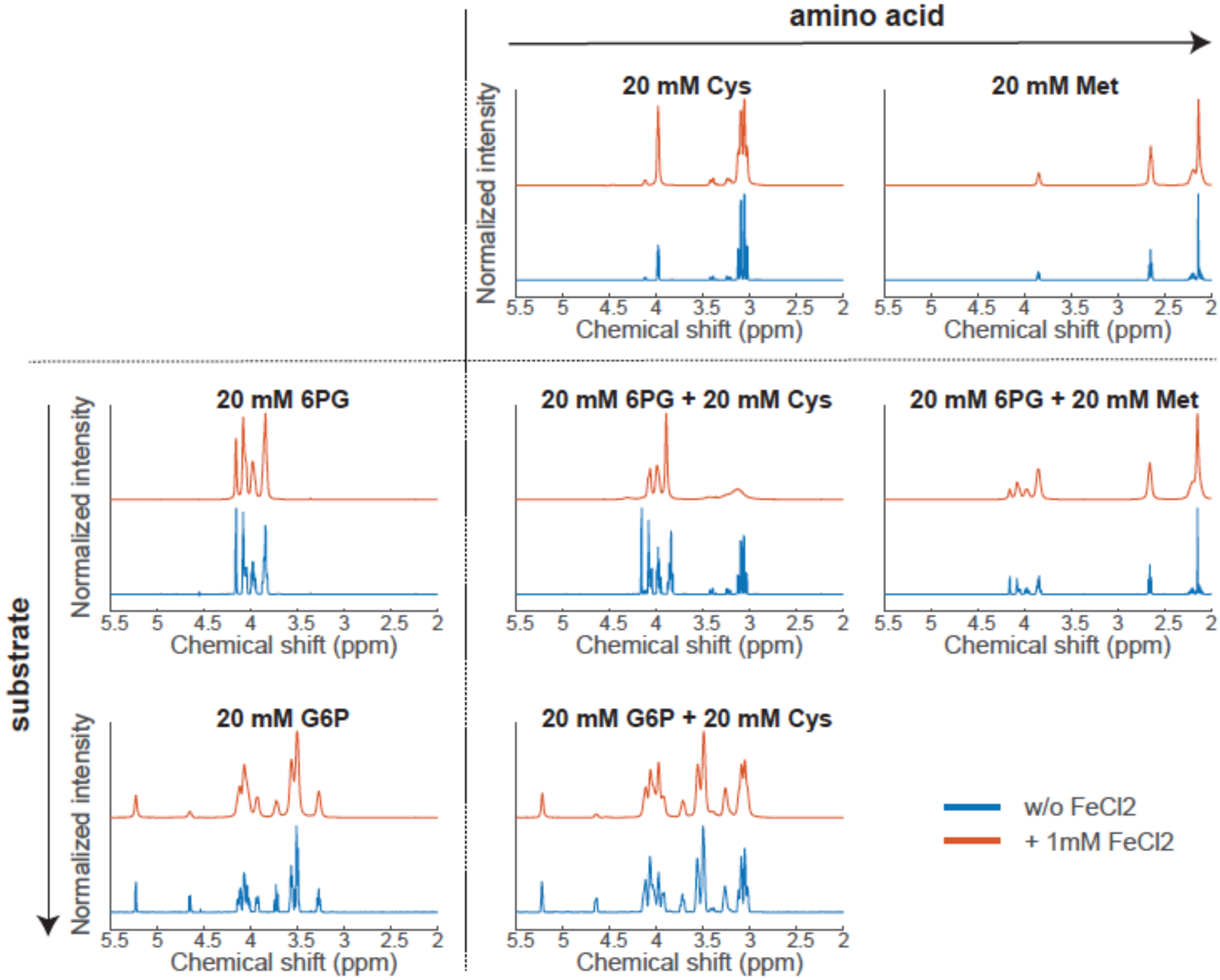


**Fig. S6. [1]H-NMR spectra of non-reactive substrate or amino acid substituents and interactions with paramagnetic $Fe^{2+}$.** $Fe^{2+}$ broadens the proton signal peaks of closely proximal molecules in solution (in this case solvent is 50 mM phosphate pH 5 in $D_2O$). $^1$H-NMR spectra of solutions containing 20 mM G6P + 20 mM cysteine or 20 mM 6PG + 20 mM methionine were compared with those of a solution with the key components of the reaction: 20 mM 6PG + 20 mM cysteine. Addition of 1 mM $FeCl_2$ (spectra in red) just slightly affected the individual species when analyzed separately, but extensively distorted the peaks of 6PG and cysteine when these were combined in the same solution, in contrast with the other two cases with either G6P or methionine where peak definition remained almost unaltered. This suggests a fairly specific cysteine-6PG interaction that makes them more likely to $Fe^{2+}$ binding. In all cases, representative examples are shown from at least $N$ = 3 independent experiments (for visual comparison, spectra are shown normalized to the maximum peak intensity in each case).

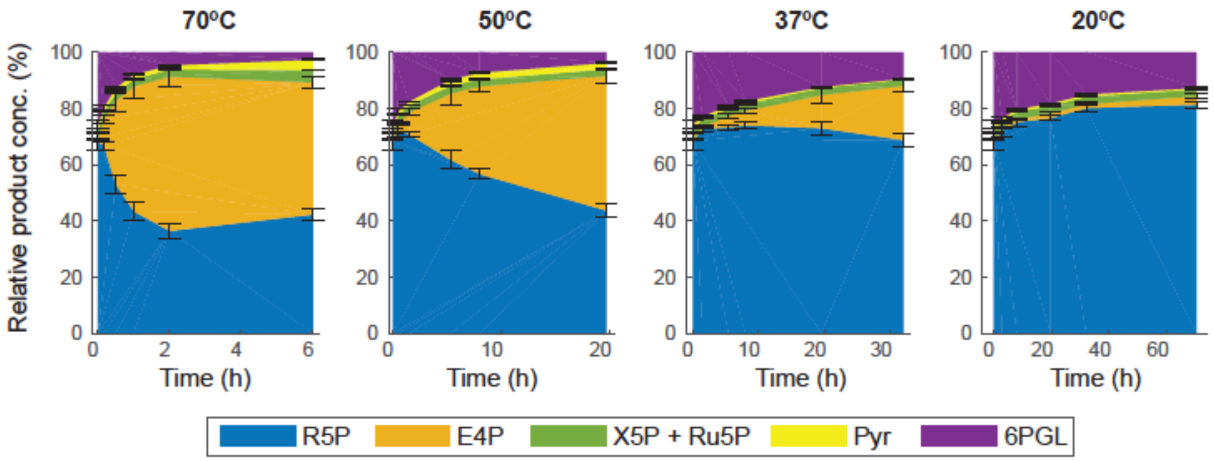


**Fig. S7. Thermal-dependent product specificity over time.** 800 µM 6PG was incubated at different temperatures in 50 mM phosphate solution pH 5 containing 75 µM cysteine and 75 µM $FeCl_2$, and the formation of different sugar-phosphate products monitored over time. Shown is the cumulative product concentration normalized to the total product quantified at each time by targeted LC/MS. As temperature decreases, the loss of specificity in R5P formation detected at higher temperatures ceases and R5P becomes the predominant sugar-phosphate product, also at long term (70-80% of total product; blue area). Time ranges are the same as in main Fig. 3A and were defined based on the kinetics scaling with temperature. Error bars account for mean ± SD from $N = 3$ independent experiments.

**Table S1. Sugar-phosphate interconversion rates at 70°C in the presence of different amino acids. See table as separate file.**

**Table S2. Effect of oxygen on the formation of ribose 5 phosphate.**

| Sl | Reagents* | Ratio of R5P concentration formed under aerobic vs. anaerobic environment |
|---|---|---|
| 1 | 400 µM 6PG, 150µM $Fe^{2+}$ and 150µM Cys | 3.05 |
| 2 | 400 µM 6PG and 150µM Cys | 1.11 |
| 3 | 400 µM 6PG and 150µM $Fe^{2+}$ | 2.5 |
| 4 | 400 µM 6PG alone | 1 |

* Reagents were dissolved in phosphate solution 50 mM pH 5.0. The reaction mixture was heated at 70°C for 3h under respective environments (ambient or anaerobic chamber). 6PG: 6-phosphogluconate, R5P: ribose 5-phosphate.

**Table S3. Measurement of trace Fe contamination in the reagents used.**

| **Component** | **Concentration ± RSD (µM)** |
|---|---|
| Cysteine | 1.065 ± 0.005 |
| 6-phosphogluconate | 0.999 ± 0.007 |
| Phosphate solution | 0.938 ± 0.011 |
| Cys+6PG under standard reaction conditions* | 1.108 ± 0.011 |

*Dissolved in 50 mM phosphate solution at pH 5.0 and incubated in amber vials at 70°C for 3h.

**Table S4. $T_1$ relaxation time of cysteine solutions with increasing concentrations of Fe(II).**

| Fe ratio | 3.97 ppm | 3.06 ppm |
|---|---|---|
| 0 | 10.82 ± 0.77 | 3.10 ± 0.17 |
| 0.005 | 6.11 ± 0.20 | 2.63 ± 0.01 |
| 0.01 | 4.54 ± 0.08 | 2.56 ± 0.03 |
| 0.05 | 3.75 ± 0.34 | 2.52 ± 0.20 |

**Table S5. $T_1$ relaxation time of 6PG solutions with increasing concentrations of Fe(II).**

| Fe ratio | 4.12 ppm | 4.09 ppm | 3.96 ppm | 3.84 ppm |
|---|---|---|---|---|
| 0 | 2.74 ± 0.13 | 2.07 ± 0.06 | 0.98 ± 0.01 | 2.23 ± 0.01 |
| 0.005 | 1.35 ± 0.05 | 1.24 ± 0.06 | 0.74 ± 0.02 | 1.31 ± 0.03 |
| 0.01 | 1.21 ± 0.05 | 1.12 ± 0.04 | 0.71 ± 0.04 | 1.19 ± 0.03 |
| 0.05 | 0.89 ± 0.06 | 0.85 ± 0.04 | 0.58 ± 0.05 | 0.97 ± 0.03 |

**Table S6. $T_1$ relaxation time of mixed cysteine-6PG solutions with increasing concentrations of Fe(II).**

| Fe ratio | 4.17 ppm | 4.08 ppm | 4.00 ppm | 3.85 ppm | 3.10 ppm |
|---|---|---|---|---|---|
| 0 | 3.24 ± 0.31 | 2.19 ± 0.09 | | 2.28 ± 0.12 | 2.51 ± 0.21 |
| 0.005 | 0.97 ± 0.03 | 1.42 ± 0.03 | 1.10 ± 0.02 | 1.30 ± 0.02 | 1.60 ± 0.02 |
| 0.01 | 0.69 ± 0.05 | 1.02 ± 0.06 | 0.74 ± 0.02 | 0.94 ± 0.05 | 1.20 ± 0.03 |
| 0.05 | | 0.46 ± 0.08 | 0.35 ± 0.04 | 0.53 ± 0.04 | 0.49 ± 0.05 |

**Table S7. $T_1$ relaxation time of methionine solutions with increasing concentrations of Fe(II).**

| Fe ratio | 3.85 ppm | 2.63 ppm | 2.16 ppm |
|---|---|---|---|
| 0 | 6.55 ± 0.10 | 2.51 ± 0.03 | 3.34 ± 0.03 |
| 0.005 | 4.65 ± 0.33 | 2.32 ± 0.05 | 3.16 ± 0.05 |
| 0.01 | 5.10 ± 0.19 | 2.36 ± 0.03 | 3.22 ± 0.04 |
| 0.05 | 6.21 ± 0.24 | 2.53 ± 0.01 | 3.48 ± 0.01 |

**Table S8. $T_1$ relaxation time of mixed methionine-6PG solutions with increasing concentrations of Fe(II).**

| Fe ratio | 4.19 ppm | 4.09 ppm | 3.96 ppm | 3.85 ppm | 2.63 ppm | 2.16 ppm |
|---|---|---|---|---|---|---|
| 0 | 2.33 ± 0.20 | 1.32 ± 0.10 | 0.88 ± 0.07 | 2.61 ± 0.06 | 2.33 ± 0.02 | 3.18 ± 0.01 |
| 0.005 | 1.18 ± 0.03 | 0.92 ± 0.02 | 0.69 ± 0.02 | 1.58 ± 0.03 | 1.94 ± 0.02 | 2.64 ± 0.02 |
| 0.01 | 1.02 ± 0.03 | 0.84 ± 0.02 | 0.65 ± 0.02 | 1.42 ± 0.02 | 1.84 ± 0.02 | 2.50 ± 0.03 |
| 0.05 | 0.89 ± 0.06 | 0.78 ± 0.03 | 0.66 ± 0.02 | 1.19 ± 0.03 | 1.63 ± 0.03 | 2.24 ± 0.05 |

**Table S9. $T_1$ relaxation time of G6P solutions with increasing concentrations of Fe(II).**

| Fe ratio | 5.22 ppm | 3.99 ppm | 3.92 ppm | 3.87 ppm | 3.70 ppm | 3.56 ppm | 3.50 ppm | 3.27 ppm |
|---|---|---|---|---|---|---|---|---|
| 0 | 4.02 ± 0.21 | 0.98 ± 0.03 | 0.93 ± 0.01 | 2.02 ± 0.06 | 4.06± 0.19 | 1.80± 0.02 | 2.70 ± 0.03 | 4.11 ± 0.08 |
| 0.005 | 3.60 ± 0.50 | 0.73 ± 0.02 | 0.73 ± 0.02 | 1.47 ± 0.08 | 3.02 ± 0.22 | 1.53 ± 0.06 | 2.10 ± 0.06 | 3.27 ± 0.14 |
| 0.01 | 3.85 ± 0.14 | 0.80 ± 0.04 | 0.78 ± 0.03 | 1.61 ± 0.06 | 3.31 ± 0.13 | 1.62 ± 0.04 | 2.20 ± 0.09 | 3.50 ± 0.14 |
| 0.05 | 4.19 ± 0.39 | 1.03 ± 0.01 | 0.92 ± 0.02 | 1.64 ± 0.10 | 3.50 ± 0.27 | 1.82 ± 0.03 | 2.42 ± 0.08 | 3.86 ± 0.16 |

**Table S10. $T_1$ relaxation time of mixed G6P-cysteine solutions with increasing concentrations of Fe(II).**

| Fe ratio | 5.22 ppm | 3.99 ppm | 3.92 ppm | 3.88 ppm | 3.85 ppm | 3.70 ppm | 3.56 ppm | 3.50 ppm | 3.27 ppm | 3.03 ppm |
|---|---|---|---|---|---|---|---|---|---|---|
| 0 | 4.08 ± 0.70 | 1.25 ± 0.09 | 0.99 ± 0.02 | 8.41 ± 0.25 | 2.56 ± 0.24 | 4.80± 0.60 | 1.89± 0.09 | 2.81 ± 0.06 | 4.41 ± 0.39 | 3.05 ± 0.13 |
| 0.005 | 4.75 ± 0.40 | 1.06 ± 0.07 | 0.87 ± 0.03 | 3.44 ± 0.01 | 2.12 ± 0.13 | 4.27 ± 0.35 | 1.83 ± 0.09 | 2.45 ± 0.07 | 3.84 ± 0.19 | 2.50 ± 0.08 |
| 0.01 | 3.34 ± 0.44 | 1.10 ± 0.22 | 0.84 ± 0.09 | 2.56 ± 0.16 | 2.02 ± 0.32 | 3.79 ± 0.50 | 1.68 ± 0.20 | 2.15 ± 0.09 | 3.31 ± 0.35 | 2.11 ± 0.12 |
| 0.05 | 3.05 ± 0.19 | 0.92 ± 0.02 | 0.75 ± 0.01 | 1.47 ± 0.03 | 1.56 ± 0.03 | 2.81 ± 0.04 | 1.56 ± 0.03 | 1.71 ± 0.05 | 2.59 ± 0.07 | 1.79 ± 0.05 |

### SI References

1. G. L. Ellman, Tissue sulfhydryl groups. *Arch. Biochem. Biophys.* **82**, 70–77 (1959).
